# The gut bacterial natural product colibactin triggers induction of latent viruses in diverse bacteria

**DOI:** 10.1101/2021.05.24.445430

**Authors:** Justin E. Silpe, Joel W. H. Wong, Siân V. Owen, Michael Baym, Emily P. Balskus

## Abstract

Colibactin is a chemically unstable small molecule genotoxin produced by multiple different bacteria, including members of the human gut microbiome.^1,2^ While the biological activity of colibactin has been extensively investigated in mammalian systems,^3^ little is known about its effects on other microorganisms. Here, we discover that colibactin targets bacteria carrying prophages, inducing lytic development via the bacterial SOS response. DNA, added exogenously, protects bacteria from colibactin, as does expressing a colibactin resistance protein (ClbS) in non-colibactin-producing cells. The prophage-inducing effects we observe apply broadly across taxonomically diverse phage-bacteria systems. Finally, we identify bacteria that possess colibactin resistance genes but lack colibactin biosynthetic genes. Many of these bacteria are infected with predicted prophages, and we show that the expression of their ClbS homologs provides immunity from colibactin-triggered induction. Our study reveals a mechanism by which colibactin production could impact microbiomes and highlights an underappreciated role for microbial natural products in influencing population-level events such as phage outbreaks.

## Introduction

Bacterial natural products display remarkable structural complexity and diversity, which contribute to their impressive breadth of biological activity. Unsurprisingly, natural products of microbial origin, or their derivatives, have been a productive source of antibiotics (e.g. penicillin, streptomycin) and anticancer agents (e.g., staurosporine, doxorubicin).^4,5^ While useful for therapeutic purposes, the clinical effects of most natural products require concentrations much higher than those produced in natural environments. The functions these natural products possess under conditions that are relevant to the producing organisms remain mysterious.^6,7^ In environments with rich microbial communities – whether terrestrial, aquatic, or within the human host – understanding the contributions of bacterially-produced small molecules to microbiome composition and function continues to be a challenging but important endeavor.

A bacterial natural product of particular relevance to human health is colibactin, a chemically reactive small molecule genotoxin produced by gut bacteria harboring a 54 kb hybrid nonribosomal peptide synthetase-polyketide synthase (NRPS-PKS) biosynthetic gene cluster known as the *pks* island (Figure 1a). This gene cluster is predominantly found in human-associated *Escherichia coli* strains belonging to phylogenetic group B2 but is also present in other human gut Enterobacteriaceae, as well as bacteria from the honey bee gut, a marine sponge, and an olive tree knot.^8–10^ Mechanistic studies have revealed that colibactin induces interstrand DNA cross-links in vitro, causes cell-cycle arrest in eukaryotic cell culture, and impacts tumor formation in mouse models of colorectal cancer (CRC).^1,11–13^ Colibactin-DNA adducts have been detected in mammalian cells and in mice,^14^ and recent studies have identified colibactin-associated mutational signatures in cancer genomes, predominantly from CRC.^15,16^ Despite its important biological activity, colibactin has eluded traditional isolation and structural elucidation. Information regarding its chemical structure has largely been derived from bioinformatic analysis of the gene cluster, characterization of key biosynthetic enzymes, stable-isotope feeding experiments, gene deletion studies, isolation of shunt products, and chemical synthesis of putative analogs.^17–19^ These studies suggest that colibactin possesses a pseudodimeric structure, with a reactive cyclopropane warhead at each end accounting for its characteristic DNA alkylating ability (Figure 1a).^14,20,21^

**Figure 1.**
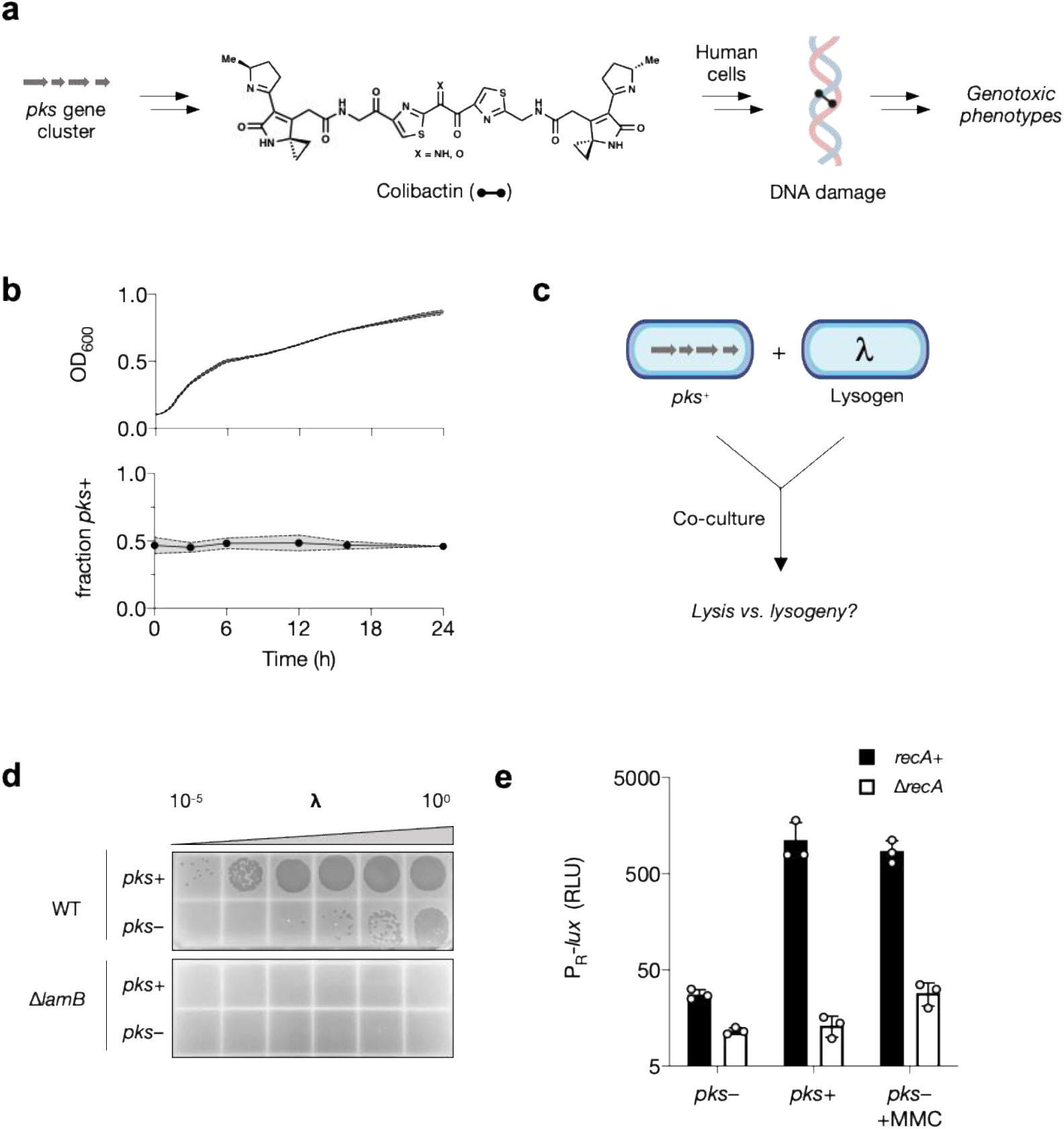
Colibactin production specifically affects prophage-carrying bacteria. **a** Structure of the genotoxin colibactin and its proposed mode of action toward mammalian cells. **b** Growth and relative abundance of *pks*^−^ and *pks*^+^ *E. coli* in co-culture. Upper, total culture density of *pks*^−^/*lacZ*^−^ *E. coli* co-cultured with *pks*^+^/*lacZ*^+^ *E. coli* at a starting ratio of 1:1; lower, the proportion of *lacZ*^+/−^ within the same co-culture based on differential blue-white plating over time (see Figure S1b for swapped markers). **c** Schematic representation of co-culture setup used throughout this work. *pks*^+^ *E. coli* is co-cultured with non-colibactin-producing *E. coli* that is infected with a temperate phage, lambda (λ). **d** Plaque assay obtained from a 24 h co-culture between *pks*^+^ or *pks*^−^ *E. coli* harboring phage lambda. Supernatants were spotted onto wild-type *E. coli* (top) and the λ-resistant Δ*lamB* mutant (bottom). **e** Relative light units (RLU) produced from a bioluminescent reporter encompassing the DNA-damage inducible region of phage lambda that regulates lysis-lysogeny (called P_R_-*lux*). Reporter output measured in *recA^+^* (black) and Δ*recA* (white) *E. coli* co-cultured with *pks*^+^ or *pks*^−^ *E. coli* in the absence or presence of 100 ng mL^−1^ MMC. Relative light units (RLU) were calculated by dividing bioluminescence by OD_600_. Data represented as mean ± SD with n = 3 biological replicates (b and e); or n = 3 biological replicates from which a single representative image is shown (d).

In contrast to its effects on higher organisms, the impacts of colibactin on the surrounding microbial community remain largely unknown. Previous studies have indicated that colibactin production may cause broad shifts in gut microbial community composition in mice and inhibit the growth of a subset of *Staphylococci*.^22,23^ However, exposure to colibactin did not affect the growth of the vast majority (97%) of bacterial species tested in this prior study, and there is no mechanistic understanding of these reported activities.

In this study, we investigated the interaction between colibactin-producing and -susceptible bacteria. We show that exposure to colibactin affects bacteria harboring prophages (latent viruses residing within the bacterium’s chromosome). Specifically, we discover that colibactin activates lytic development in prophage-carrying bacteria and that this effect occurs via the canonical SOS response. Notably, we observe that colibactin is an effective inducer of prophages across diverse phage-host systems. We also find many examples of non-colibactin-producing bacteria that possess colibactin resistance genes, indicating past exposure to this genotoxin in microbiomes. Though many of these bacteria are uncharacterized, in the cases we tested, expression of these predicted resistance genes conferred protection from prophage induction by colibactin. We propose that the occurrence of colibactin resistance in these non-producing organisms serves as a unique defense mechanism to avoid lysis and to prevent phage outbreaks caused by colibactin producers within a given community. Altogether, these discoveries highlight a role for bacterial natural products in mediating the activity of phages in microbiomes.

## Results

Our knowledge surrounding the biological activity of colibactin has come from studies in mammalian cell systems in which colibactin-producing bacteria induce hallmark genotoxic effects, such as premature senescence and cell-cycle arrest, whereas supernatants or lysates from these bacteria do not.^1,24^ Proposed explanations for why live bacteria must be present in these assays include that colibactin needs to be produced continuously due to its instability and that delivery into mammalian cells may be contact-dependent.^1^ As there are no known mechanisms by which colibactin acts beyond eukaryotic cells, we aimed to shed additional light on colibactin’s activity and potential ecological roles in microbiomes by studying its effects on bacteria.

We hypothesized that exposure to colibactin might impair bacterial cell division, and we sought to test this proposal by monitoring cell growth. To begin, we exposed a laboratory strain of non-colibactin producing (*pks*^−^) *E. coli* (BW25113) to supernatants from overnight cultures of colibactin-producing *E. coli* (a heterologous expression strain harboring the *pks* gene cluster encoded on a bacterial artificial chromosome, BAC-*pks*, hereafter called *pks*^+^). Culture supernatants did not inhibit the growth of the laboratory *E. coli* strain, matching previous reports with mammalian cells (Extended Data Figure 1a). To test if bacterial growth inhibition requires the presence of live colibactin-producing cells, we co-cultured *pks^+^ E. coli* with *pks*^−^ *E. coli* carrying chromosomally-distinguishable markers (*lacZ*) and monitored the growth of the two populations using *lacZ* as a proxy for *pks*. Figure 1b shows that, when started at a 1:1 ratio, the proportion of *pks^+^ E. coli* did not change over the course of the experiment, and this outcome occurred irrespective of which strain carried the *lacZ* marker (Extended Data Figure 1b). These results suggest that, at least under the conditions tested, colibactin production by one bacterium does not inhibit growth of an isogenic, non-producing strain.

Given its well-characterized ability to induce DNA damage and cell-cycle arrest in mammalian cells,^1^ we were struck by the lack of competitive growth advantage associated with colibactin production in bacterial co-culture. Multiple lines of evidence suggest that bacteria should be susceptible to colibactin-mediated DNA damage. For example, the final gene in the *pks* gene cluster, *clbS*, encodes a self-resistance protein reported to hydrolyze and destroy the reactive cyclopropane warheads of colibactin,^25,26^ while another gene, *clbP*, encodes a periplasmic peptidase that converts an inactive late-stage biosynthetic intermediate (precolibactin) to the final genotoxic metabolite in the periplasm before export.^27,28^ Both bacterially encoded self-resistance mechanisms suggest that, like many toxic bacterial natural products, colibactin is potentially deleterious to non-producing bacteria that lack these resistance strategies.

With the above findings in mind, we considered alternative consequences of colibactin-mediated DNA damage beyond inhibition of bacterial growth. One potential response of interest is phage induction. Specifically, it is known that DNA damage induced by UV irradiation or chemical treatment (e.g. mitomycin C, MMC) activates lytic replication of prophages (a latent form of phage infection) in bacteria, which could subsequently propagate the infection throughout the larger microbial community.^29^ We therefore wondered whether colibactin could impact bacterial populations by activating resident phages.

To test if colibactin production alters prophage behavior in neighboring, non-colibactin-producing lysogens, we infected WT *E. coli* BW25113 with phage lambda and co-cultured this lysogen with *pks*^+^ or *pks*^−^ *E. coli* (Figure 1c). As shown in Figure 1d, 24 h co-culture with the *pks*^+^ strain increased phage titers 1,000-fold above those obtained with the *pks*^−^ strain. Consistent with the resident prophage being the responsible agent, no plaques were observed from any condition on Δ*lamB E. coli*, which lacks the lambda phage receptor (Figure 1d). These results suggest that colibactin specifically affects prohage-carrying bacteria by inducing lytic development.

Regulation of lambda induction from the prophage state (the lysis-lysogeny decision) occurs via a repressor protein that blocks the launch of the phage’s lytic program. The lambda repressor protein (cI) is inactivated by the host-encoded SOS response for which RecA is a master regulator.^30^ To test if prophage induction by colibactin depends on a similar sequence of events, we engineered a transcriptional reporter to track the lambda lysis-lysogeny fate decision by fusing the lambda immunity region to the luciferase operon (*lux*) on a plasmid (hereafter called P_R_-*lux*). Light production by luciferase therefore reports the transcriptional de-repression of phage lambda lytic replication. Consistent with what is known about phage lambda, reporter-driven light production was low in WT *E. coli* but induced highly upon addition of the known DNA-damaging-, prophage-inducing agent, MMC (Extended Data Figure 1c). To examine the effect of colibactin in this system, we co-cultured *E. coli* harboring P_R_-*lux* with *pks*^+^ *or pks*^−^ *E. coli*. As expected, the *pks*^−^ strain did not activate reporter-driven P_R_-*lux* activity, nor did it prevent the reporter strain from being induced with MMC (Figure 1e). In contrast, co-culture with the *pks*^+^ strain induced P_R_-*lux* in the reporter strain 40-fold (Figure 1e). Notably, the activating effect of both MMC and co-cultured *pks*^+^ cells was eliminated when the P_R_-*lux* plasmid was transformed in Δ*recA* versus *recA*^+^ *E. coli* (Figure 1e) showing that the transcriptional de-repression requires the SOS response. These results, taken together, suggest that colibactin causes prophage induction, and therefore cell lysis, of neighboring *pks*^−^ bacteria via the canonical DNA-damage inducible SOS pathway.

We next sought to further pinpoint the factors responsible for the observed phage-inducing activity of colibactin. During colibactin biosynthesis, activation of the precolibactin ‘prodrug’ by the periplasmic peptidase ClbP generates the reactive cyclopropane warheads essential for genotoxicity (Figure 2a).^27,28,31^ Consistent with these findings, deletion of *clbP* in the producing strain abolished activation of P_R_-*lux* activity in co-cultured reporter cells (Figure 2b). As depicted in Figure 2a, when the same *clbP* deletion strain was co-cultured with the lambda lysogen, we observed a significant reduction in phage titers compared to the *pks*^+^ strain (Figure 2c). Another important consideration is that many studies of colibactin employ strains that heterologously express the *pks* cluster on a BAC.^32^ This strategy has proven useful for elucidating colibactin’s structure and biosynthesis, but it may fail to capture regulatory features of native colibactin-producing strains. To address this uncertainty, we tested a native colibactin producer, *E. coli* NC101, a murine adherent-invasive strain isolated from mouse models of CRC carcinogenesis.^13,33^ As shown in Figure 2b and c, WT NC101 activated the P_R_-*lux* reporter and increased phage titers by approximately 2 orders of magnitude compared to the NC101 Δ*clbP* mutant. While the number of plaques produced was generally lower from the NC101 co-cultures than the heterologously expressed BAC-*pks*, the result demonstrates that a native colibactin-producer induces the SOS response and activates prophages in neighboring cells in a *pks*-dependent manner, and in comparable magnitude compared to the BAC-*pks*-containing laboratory *E. coli*. Moreover, our reporter data showing that PR-*lux* activity is completely abolished when exposed to the NC101 Δ*clbP* mutant, suggests that, under the conditions tested, colibactin production is likely the only mechanism by which NC101 activates the SOS response in neighboring *E. coli*.

**Figure 2.**
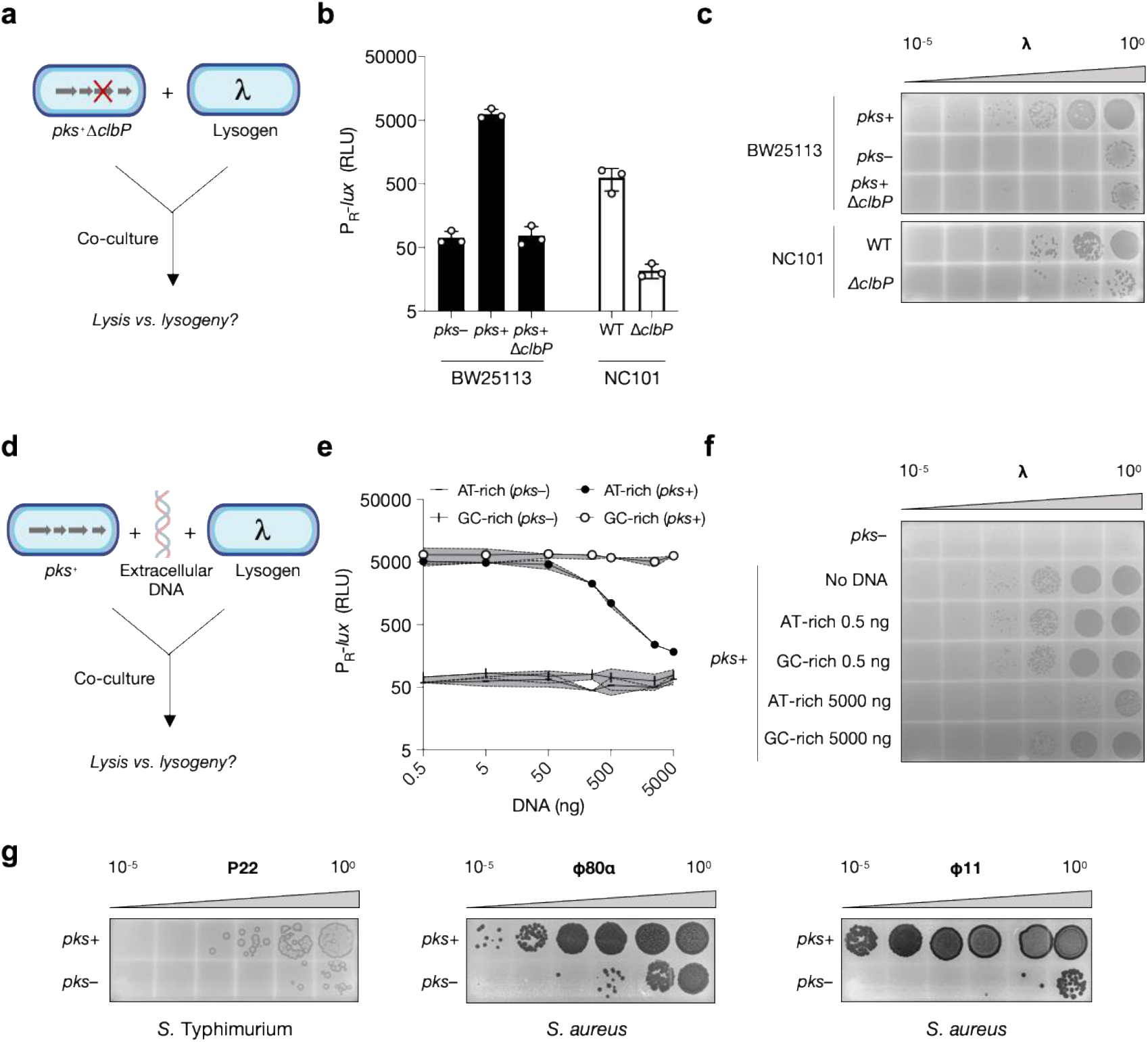
Prophage induction is dependent on colibactin-mediated DNA alkylation and affects diverse phage-bacteria systems. **a** Schematic of co-culture experiment with a colibactin biosynthesis-defective (Δ*clbP*) *pks* strain. **b** P_R_-*lux* output from reporter cells co-cultured with *E. coli* BW25113 (*pks*^+^, *pks*^−^, and *pks*^+^Δ*clbP;* black bars) or native-colibactin producing *E. coli*, NC101 (WT and Δ*clbP*; white bars). **c** Plaque assays obtained from analogous incubations as in (a) but with a lambda lysogen used in place of the reporter strain. **d** Schematic of co-culture experiment in which extracellular DNA (eDNA) is added to the media. **e** P_R_-*lux* output from reporter cells co-cultured with *pks*^+^ *E. coli* in the presence of varying amounts of extracellular DNA (AT-rich and GC-rich DNA, black and white symbols, respectively). **f** Plaque assays of the analogous incubations as in (c) but with a lambda lysogen used in place of the reporter strain. **g** Plaque assays for each of the lysogens (bacterium / phage) indicated after co-culture with *pks*^+^ or *pks*^−^ *E. coli*. In b and e, RLU as in 1e. Data represented as mean ± SD with n = 3 biological replicates (b and e); or n = 3 biological replicates from which a single representative image is shown (c, f, and g).

To further interrogate the response of neighboring bacteria to colibactin production, we sought to manipulate the amount of genotoxin delivered to target organisms (Figure 2d). In mammalian cell systems, addition of extracellular DNA has been shown to attenuate the genotoxicity of *pks*^+^ *E. coli*, presumably by titrating away the reactive colibactin through alkylation of the exogenous DNA.^11^ Consistent with this result, and supporting the model that colibactin is released extracellularly, addition of herring sperm DNA to our bacterial co-culture system resulted in a DNA concentration-dependent reduction in P_R_-*lux* activity (Extended Data Figure 2a) and phage titers (Extended Data Figure 2b). To further investigate this effect, we designed short oligonucleotides with variable AT:GC ratios relative to herring sperm DNA (58% AT). AT-rich DNA (75% AT) but not GC-rich DNA (29% AT) attenuated prohage-induction, both in terms of reporter output and plaques produced (Figures 2e and f). The pattern we observe for AT-versus GC-rich DNA in our assay is consistent with recent reports of colibactin-induced DNA damage and its concomitant mutational signatures occurring predominantly at AT-rich motifs.^15,16^ Taken together, our observations that colibactin-producing bacteria require ClbP to induce prophages in nearby cells and that exogenous addition of DNA ameliorates this effect, strongly suggest that, as has been shown in eukaryotic systems, the ability to produce and transmit the final genotoxic product is important for colibactin’s effect on bacteria.

Given that bacteria frequently exist in polymicrobial communities (microbiomes), that prophages are pervasive in these communities, and that the SOS response is highly conserved, we wondered if the genotoxic effect of colibactin could also induce prophages residing in phylogenetically distinct, gram-negative and -positive bacteria. To explore this, we co-cultured *pks*^+^ and *pks*^−^ *E. coli* with prophage-carrying *Salmonella enterica* serovar Typhimurium (harboring prophage P22) and *Staphylococcus aureus* (harboring prophages phi11 and phi80α). All three phage-bacteria systems showed a *pks*-dependent increase in plaques produced (Figure 2g). Because cell lysis is an irreversible consequence of prophage induction,^29^ we wondered whether prophage-mediated cell lysis could explain a previously reported observation of *pks*-dependent growth inhibition of a subset of *S. aureus* strains.^23^ We found that when co-cultured with *pks^+^ E. coli*, the two prophage-carrying *S. aureus* strains tested underwent a four order of magnitude drop in colony forming units (CFU) when cultured with *pks*^+^ versus *pks*^−^ *E. coli*, as compared to a two order of magnitude drop observed for phage-free *S. aureus* (Extended Data Figure 3). These results suggest that prophage-mediated cell lysis enhances the ability of colibactin-producing bacteria to outcompete susceptible members of a mixed population.

These results predict that colibactin, like MMC, is a generally effective inducer of prophages. However, unlike MMC, where self-protection to the producing organism is thought to require the combined action of several different resistance proteins,^34,35^ protection from colibactin in *pks*^+^ organisms involves one 170 amino acid resistance protein, ClbS.^25,26^ Acquisition of ClbS would therefore be a potential strategy for susceptible community members to acquire resistance against the effects of colibactin. We thus wondered whether *clbS*-like genes might exist in closely related, non-colibactin producing bacteria, and whether the function of these genes may alter phage-host dynamics in response to colibactin. To first gain insight into its context, we performed a bioinformatic search (tBLASTn) for genes that translate to products precisely matching the *E. coli* ClbS amino acid sequence. In 97% of the examined hits (230 total, Supplementary Table 1), the *clbS* homolog was found in a *pks* gene cluster having the same genetic organization as known colibactin-producing strains. In the 7 cases (3%) where *clbS* was not associated with an intact *pks* gene cluster, the gene normally encoded upstream of *clbS*, *clbQ*, was present but truncated, and both genes were surrounded by predicted transposases or transposase-associated genes (Extended Data Figure 5a and Supplementary Table 1), indicating the region may be subject to horizontal transfer. This search, consistent with a recent report on ClbS-like proteins,^36^ reveals that the *clbS* gene found in *pks*^+^ *E. coli* also exists in isolates of the same species lacking a *pks* gene cluster.

To test whether expression of ClbS in non-colibactin producing *E. coli* can provide protection from colibactin exposure, we introduced and expressed plasmid-encoded *clbS* (originating from *pks*^+^ *E. coli*, pTrc-*clbS*) in a *pks*^−^ strain harboring either the P_R_-*lux* reporter or prophage lambda (depicted in Figure 3a). When co-cultured with *pks*^+^ *E. coli*, the reporter strain harboring the *clbS* expression vector repressed P_R_-*lux* reporter activity whereas the same reporter strain transformed with pTrc-Δ*clbS* (a vector isogenic to pTrc-*clbS*, lacking all except the first and last 24 nt of *clbS*) did not (Figure 3b). ClbS did not prevent P_R_-*lux* reporter activity when MMC was used as the inducing agent (Extended Data Figure 4), suggesting that this protection is specific to colibactin. Consistent with the reporter activity, the lambda lysogen harboring pTrc-*clbS* yielded 1,000-fold fewer phage particles than the same lysogen carrying pTrc-Δ*clbS* when in co-culture with *pks^+^ E. coli* (Figure 3c).

**Figure 3.**
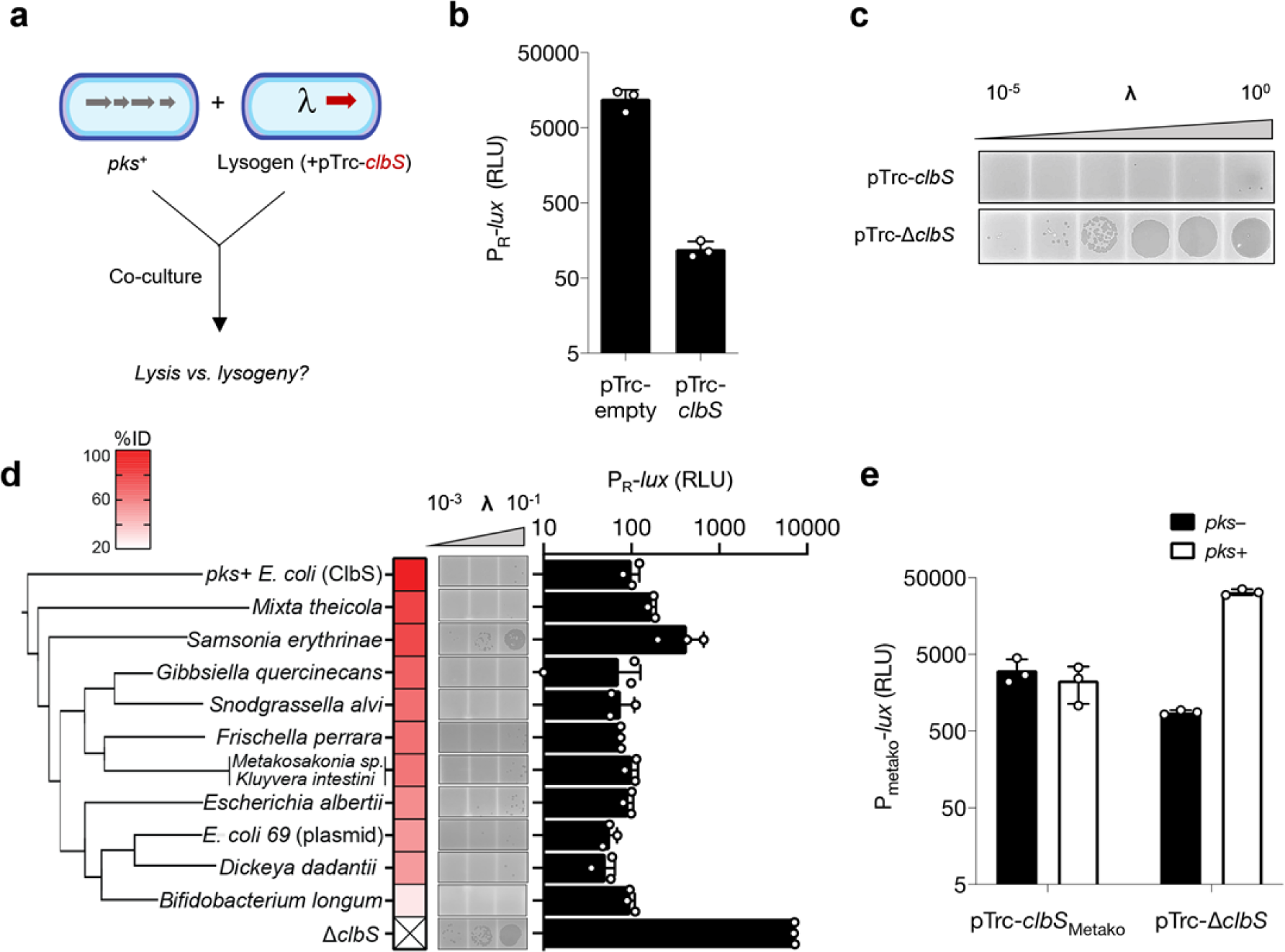
ClbS and ClbS-like proteins from diverse bacteria provide protection against colibactin-activated prophage induction. **a** Schematic of co-culture experiment with the gene encoding colibactin resistance, *clbS*, expressed *in trans*. **b** P_R_-*lux* reporter output obtained from *pks*^+^ *E. coli* co-cultured with *pks*^−^ *E. coli* harboring the reporter plasmid and either pTrc-*clbS* or the same vector with *clbS* removed (pTrc-Δ*clbS*) expressed *in trans*. **c** Plaque assays obtained from analogous incubations as in (b) but with a lambda lysogen used in place of the reporter strain. **d** Plaque assay (micrographs, left) and P_R_-*lux* reporter output (bar graph, right) obtained from *pks^+^ E. coli* co-cultured with *pks*^−^ *E. coli* harboring a vector encoding the *clbS*-like gene of the indicated organism or the pTrc-Δ*clbS* vector. For plaque assays, the *pks*^−^ *E. coli* strain harbored phage lambda, for the P_R_-*lux* reporter assay, the *pks*^−^ *E. coli* strain harbored the reporter plasmid. Heatmap and clustering of the ClbS-like proteins are based on amino acid identity to *E. coli* CFT073 ClbS. *Metakosakonia sp*. and *K. intestini* share the same ClbS amino acid sequence **e** P_Metako_-*lux* reporter output from co-cultures of *pks*^−^ or *pks*^+^ *E. coli* with *E. coli* harboring the P_Metako_-*lux* plasmid and a second plasmid (pTrc-*clbS_Metako_* or pTrc-Δ*clbS*), as indicated on the x-axis. In c and d (plaque assays), data shown as a single representative image from n = 3 biological replicates; for b, d, and e (*lux* reporter) data represented as mean ± SD with n = 3 biological replicates and RLU as in 1e.

Having confirmed that ClbS protects prophage-carrying *E. coli* from colibactin-induced lytic development, we next sought to examine whether *clbS*-like genes exist in more distantly related bacteria that lack the *pks* gene cluster; and, if so, whether they also serve a protective function in this context. Based on the assumption that organisms found in close proximity with colibactin producers would benefit from colibactin resistance, we searched for more distinct ClbS homologs (>50% amino acid identity) in the genomes of bacteria from two specific niches: the human gastrointestinal tract, encompassing known *pks* encoders such as *E. coli, Klebsiella pneumoniae*, and *Citrobacter koseri;* and the honey bee gut, which includes the colibactin-producing symbiont *Frischella perrara*.^9,37^ We found several *clbS*-like genes in *pks^−^* human-associated bacteria, including *Escherichia albertii, Kluyvera intestini* and *Metakosakonia* sp. MRY16-398, the latter two of which were isolated from patients with gastric cancer and diverticulitis, respectively.^38,39^ We also identified an additional *clbS*-like gene in *Snodgrasella alvi* wkB2, a *pks^−^* core member of the honey bee gut (Extended Data Figure 5b).^40^

To assess if these orthologs could protect against colibactin-induced phage lysis, we heterologously expressed each ClbS-like protein in the *E. coli*-lambda lysogen and co-cultured it with *pks^+^ E. coli*. As shown in Figure 3d, all 4 ClbS-like proteins attenuated prophage induction, both in terms of reporter output and plaques produced, suggesting the potential for the bacteria harboring these genes to be protected from colibactin. Removing the niche-specific criteria and lowering the cutoff in our search led us to uncover a wider range of ClbS-like proteins (25-80% amino acid identity relative to *E. coli* ClbS), an additional 6 of which we chose as a representative panel for heterologous expression in our assays (Extended Data Figure 5b). Each of these ClbS-like proteins provided protection against colibactin-induced DNA damage and prophage induction (Figure 3d). Collectively, these results demonstrate that protection from colibactin can be gained via distantly related ClbS-like proteins found in bacteria lacking all other *pks* genes.

With the above results in mind, we reasoned that protection against colibactin-mediated DNA damage could provide a mechanism by which these diverse bacteria can avoid induction of native prophages they might carry. Thus, we searched for resident prophages in the available genomic data of the 12 bacteria from which the ClbS panel was assembled. Using a phage-prediction algorithm (PHASTER),^41^ we found a total of 94 prophage regions in the 12 genomes (Extended Data Figure 6a and b), 9 of which were further classified as containing at least one intact prophage (score >90, Extended Data Figure 6b). Like their bacterial host, these putative phages are largely uncharacterized, however, we hypothesize based on domain analysis of the predicted prophage repressors that a subset of these phages are responsive to DNA damage in a manner similar to phage lambda (Extended Data Figure 6c).

Along these lines, we aimed to test whether we could recapitulate and measure the DNA damage response from any of the predicted complete prophages residing in our panel of *clbS-* carrying, *pks^−^* bacteria. Specifically, one of the intact prophages predicted by PHASTER corresponds to a 40 kb element present in the human-associated bacterium *Metakosakonia* sp. MRY16-398 (Extended Data Figure 6c). Apart from its initial isolation, *Metakosakonia* sp. MRY16-398 is uncharacterized.^38^ Likewise, the genome of the resident 40 kb prophage lacks any obvious nucleotide or amino acid-level identity to phage lambda; however, it contains a similar immunity region, as predicted by the DNA-damage sensitive repressor (Extended Data Figure 6c). We synthesized the ~1 kb region from the *Metakosakonia* sp. MRY16-398 genome encompassing the putative *cI*-like repressor and fused the counter-oriented promoter to *lux* on a plasmid (called PMetako-*lux*). When transformed into *recA*^+^ *E. coli*, PMetako-*lux* activity was activated by MMC (Extended Data Figure 6d). Similarly, co-incubation of *E. coli* harboring the PMetako-*lux* reporter plasmid with *pks^+^* but not *pks^−^ E. coli* induced PMetako-*lux* expression (Figure 3e). These two findings corroborate the domain analysis and indicate that, analogous to the immunity region of phage lambda, this prophage in *Metakosakonia* sp. MRY16-398 displays DNA-damage inducibility.

Finally, to recapitulate both colibactin-resistance and phage regulation from *Metakosakonia sp*. MRY16-398, we tested whether the ClbS natively encoded by this organism (*clbSMetako*) would confer protection from colibactin in the *Metakosakonia*-phage-based reporter system. As shown in Figure 3e, introduction of *clbSMetako* to cells harboring the PMetako-*lux* plasmid prevented *pks*-induced activation of the reporter. These results strongly suggest that this *Metakosakonia* strain is resistant to the prophage-inducing effects of colibactin and imply that acquisition of orphan *clbS* genes, perhaps by horizontal gene transfer, might be an effective strategy for prophage-carrying bacteria to resist colibactin production by neighboring community members.

## Discussion

In this study, we investigated the effects of the genotoxic natural product colibactin on bacteria. We discovered that exposure to *pks^+^ E. coli* induces prophages of diverse phage-bacteria systems. Prophage induction by colibactin occurs via the bacterial host SOS response, indicating that colibactin, in addition to damaging DNA in mammalian systems, also damages the DNA of surrounding bacteria. Finally, we find colibactin resistance genes in bacteria that themselves do not produce colibactin, suggesting a past history of exposure and adaptation to this natural product and a previously unappreciated role for colibactin in the microbial world.

The knowledge that colibactin induces prophages in diverse bacteria, combined with the finding that non-colibactin producing bacteria from distinct environmental origins harbor functional *clbS*-like genes, leads us to speculate that colibactin production is more widespread than currently recognized, and that this genotoxin likely evolved to target bacteria rather than a mammalian host. To date, colibactin has primarily been investigated in the context of the human gut in relation to host carcinogenesis, but this carcinogenic activity raises important questions regarding the evolutionary role of this toxin for the producing bacterium. Colibactin genes have also been implicated in siderophore synthesis and microcin export, suggesting that these factors may collectively be involved in bacterial competition.^42,43^ While other functions of colibactin may exist, our discovery that it induces prophages provides one mechanism by which production of and immunity to this natural product might confer a competitive advantage over other members in a microbial community. In particular, the drastic *pks*-dependent reduction in CFUs we observe for lysogenic *S. aureus*, an organism with an evolutionary history checkered by phage infection,^44,45^ may help to explain the mechanism behind a previously reported pattern of growth inhibition upon exposure to colibactin,^23^ and lends support to the hypothesis that prophage carriage is at least one factor that predicts colibactin’s impact on microbes. The broad-spectrum activity of colibactin in inducing prophages across phylogenetically distinct bacteria is especially striking and suggests this natural product could have effects on many members of a community, potentially accounting for colibactin-associated changes in gut microbiome composition previously observed in animal models.^22^ Furthermore, because the virulence behaviors of certain pathogenic bacteria depend on their lysogenic state, it is interesting to consider how colibactin might operate in these situations either positively, by helping to eliminate the pathogen, or negatively, by activating the expression of prophage-controlled virulence factors the pathogen harbors. Such is the case in Shiga toxin (Stx) producing *E. coli*, where the use of quinolone antibiotics is specifically contraindicated because it increases the induction of the SOS-responsive Stx prophage, which carries the Stx toxin.^46^

It is also interesting to compare how phage-mediated lysis compares to more direct means of cell-killing and competition (e.g. production of bacteria-encoded toxins). For example, if there were members of the bacterial community susceptible to phages induced by colibactin, production could indirectly stimulate a phage outbreak. By inducing phage outbreaks in microbial communities, colibactin could impact bacteria beyond those which are directly exposed to the metabolite itself. Furthermore, phage infections that lead to lysogeny could have lasting consequences on the infected population as the prophage will be inherited by future generations. We also envision scenarios where prophage induction could be counterproductive for colibactin producers. For example, in a community of closely related species or strains of the same species, phage induction may create phage particles that can go on to infect the colibactin producer, if susceptible. Nevertheless, in most microbial habitats where phages are thought to significantly outnumber the bacterial population,^47,48^ it is likely the colibactin producer itself is also a lysogen and hence immune from superinfection. Such a scenario has been examined in a prophage-containing clinical isolate of *Enterococcus faecalis*, where the presence of the resident prophage enhances this isolate’s colonization over other closely related phage-susceptible strains both in vitro and in vivo.^49^

Our study also highlights major gaps in our understanding of the molecular mechanisms underlying prophage induction in microbiomes. MMC and ultraviolet light are the most common methods of activating phage lysis in the laboratory, however, the ecologically relevant triggers for prophages found in natural environments remain largely unidentified. Recent work has shown that the human gut commensal bacterium, *Lactobacillus reuteri*, harbors a prophage that undergoes induction during gastrointestinal transit.^50^ The precise mechanism for induction has not been defined, but it is known to depend on bacterial metabolism of dietary fructose and SCFAs, both of which can be found at high levels in the gastrointestinal tract.^51,52^ In the vaginal community, metabolic conversion of a tobacco smoke constituent, benzo[a]pyrene, to benzo[a]pyrenediol epoxide and subsequent secretion in the vagina promotes prophage induction in multiple *Lactobacillus* strains.^53^ In the nasal microbial community, species-specific production and susceptibility to hydrogen peroxide has been shown to selectively remove prophage-carrying susceptible bacteria.^54^ Unlike benzo[*a*]pyrene, which humans encounter via outside exposures, and hydrogen peroxide, which has a wide range of biological targets (proteins, lipids, and nucleic acids) and proposed functions (immunomodulatory, antibacterial), colibactin is a complex, small molecule natural product produced by human gut bacteria. By uncovering the phage inducing activity of colibactin-producing bacteria, our findings illustrate a previously unrecognized mechanism by which colibactin and potentially other DNA-damaging natural products may function to shape microbial communities.

More generally, the modulation of phage behaviors represents a distinct and underappreciated ecological role for microbial natural products. Recent studies have identified secondary metabolites produced by *Streptomyces*, including known chemotherapuetics (anthracyclines) and antibiotics (aminoglycosides), as inhibitors of phage infection.^55–57^ At the same time, metabolites of other bacteria, including known virulence factors (pyocyanin) and bacteria-specific signaling molecules (autoinducers), promote prophage induction.^58,59^ Our findings on colibactin, as a gut metabolite, adds to this growing understanding of how small molecules can influence phage behaviors, and importantly, demonstrates phage induction by a bacterial natural product in co-culture.

While our understanding of the connections between the gut microbiome and human health continues to strengthen, the contributions of the viral component in this community both to the resident bacteria and human host remain widely unknown. In particular, the role of phages, despite making up the vast majority of the gut virome,^60^ is poorly understood. Recent improvements in sequencing platforms and library preparation methods have allowed for a more comprehensive view of the gut virome as it relates to diseases such as type I and II diabetes,^61–63^ hypertension,^64^ IBD,^65^ and CRC.^66^ Specifically, increased temperate phage abundance, through prophage induction, has been identified as a major contributing factor to shifts in the overall virome composition in IBD.^67^ Our findings set the stage for further investigations of how gut bacterial metabolite production modulates phage behaviors and influences human disease.

## Methods

### Bacterial strains, plasmids, and routine cultivation

Bacterial strains and plasmids used in these studies are listed in Supplementary Tables 2 and 3, respectively. Unless otherwise noted, *E. coli* DH10B (NEB) was used for all strain construction and propagated aerobically in Luria-Bertani (LB-Lennox, RPI) broth at 37 °C. Oligonucleotides (Sigma) and dsDNA gene blocks (IDT) used in plasmid construction are listed in Supplementary Table 4. Plasmid construction was carried out using enzymes obtained from NEB (NEBuilder HiFi DNA assembly master mix, T4 DNA ligase, and DpnI). Growth, reporter, and lysis assays were all carried out in M9 medium supplemented with 0.4% casamino acids (M9-CAS, Quality Biological) unless otherwise specified. Antibiotics, inducers, and indicators were used at the following concentrations: 100 μg mL^−1^ ampicillin (Amp, IBI Scientific), 50 μg mL^−1^ kanamycin (Kan, VWR), 25 μg mL^−1^ chloramphenicol (Cm, Sigma), 100 ng mL^−1^ mitomycin C (MMC, Sigma), 40 μg mL^−1^ 5-Bromo-4-Chloro-3-Indolyl-beta-D-Galactosidase (X-gal, Takara Bio), and 500 μM isopropyl beta-D-1-thiogalactopyranoside (IPTG, Teknova), unless otherwise specified.

### Growth and competition assays

#### For growth inhibition by cell-free fluids

Overnight cultures of WT *E. coli* BW25113 harboring either BAC-*pks* or the empty BAC were centrifuged (16,100 g and 1 min) and the supernatant was passed through a 0.22 μm filter (Corning Spin-X). Growth of non-colibactin producing *E. coli* cultures was assayed in fresh LB in the presence of varying amounts of each supernatant (5%, 10%, 20%, 50% v/v). OD_600_ was measured at regular intervals using a BioTek Synergy HTX multi-mode plate-reader.

#### For E. coli-E. coli competition assays

Overnight cultures of *lacZ^+^ E. coli* MG1655 (KIlacZ, Addgene: #52696) harboring BAC-*pks* was back-diluted 1:100 into fresh M9-CAS and mixed in a 1:1 ratio with a similarly back-diluted culture of *lacZ*^−^ *E. coli* MG1655 (delta-Z, Addgene: #52706) harboring the empty-BAC. The co-cultures were incubated at 37 °C, and, at regular intervals, an aliquot was taken for differential plating on LB supplemented with X-gal and IPTG. Both BAC combinations (*pks^+^* versus empty) and marker combinations (*lacZ*^+^ *versus lacZ*^−^) were tested to rule out the influence of carrying the *lacZ* marker.

#### For E. coli-S. aureus competition assays

*S. aureus* RN10359 Φ80α and *S. aureus* RN10359 Φ11 were grown overnight at 37 °C in fresh Brain Heart Infusion (BHI) media, while *E. coli* BW25113 harboring BAC-*pks* or empty-BAC were grown overnight at 37 °C in fresh LB broth supplemented with Cm. The overnight cultures were back-diluted 1:100 into fresh BHI medium and mixed in a 1:1 ratio and incubated at 37 °C for 24 h. The cultures were plated on LB agar supplemented with Cm for *E. coli* CFUs, and mannitol salt phenol-red agar (Sigma) for *S. aureus* CFUs.

### Production and isolation of phage lambda by mitomycin C induction

An overnight culture of the lambda lysogen was back-diluted 1:100 into fresh LB and incubated at 37 °C. Upon reaching OD_600_ = 0.4-0.5, MMC (500 ng mL^−1^ final concentration) was added and the cultures were returned to 37 °C for an additional 3-5 h, over which time noticeable clearing occurred. After chloroform treatment and centrifugation (16,100 g and 1 min), the clarified lysates were filter-sterilized and stored at 4 °C prior to use.

### Quantification of phage induction by colibactin

#### Reporter assay

Overnight cultures were back-diluted 1:100 into fresh M9-CAS medium with appropriate antibiotics prior to being dispensed (200 μL) into white-walled 96-well plates (Corning 3610). For co-culture experiments, the two cultures were mixed 1:1 immediately after back-dilution. Monoculture controls for each strain were prepared by adding 100 μL of the back-diluted cultures to an equivalent volume of M9-CAS. For DNA interference experiments, herring sperm DNA (Promega) was used. To test DNA with varying AT richness, complementary oligonucleotide pairs [JWO-1046 and JWO-1047] and [JWO-1044 and JWO-1045] were annealed in 10 mM aqueous Tris-HCl buffer, and the resulting duplexes were added to the wells at the indicated concentrations. Plates were shaken at 37 °C and the OD_600_ and bioluminescence readings were obtained after 24 h. Relative light units (RLU) were calculated by dividing the bioluminescence by the OD_600_.

#### Plaque assays

Preparing and measuring viral titers from phage-infected co-cultures was carried out according to the identical conditions used for the reporter assays with the exception that the reporter strain was substituted for the lambda lysogen. To prepare phage lysates, cultures were transferred after overnight growth to microcentrifuge tubes and centrifuged at 16,100 g for 1 min). The supernatant was removed and passed through a 0.22 μm filter. Supernatants were diluted logarithmically from 10^0^ to 10^−5^), and 10 μL spotted on top-agar (preparation below) containing the lambda-sensitive indicator strain (WT BW25113) or the resistant control (*lamB::kan*) *E. coli*.

#### Preparation of top agar

Overnight cultures of WT BW25113 or the *lamB::kan E. coli* (for lambda phage enumeration); phage-free strain *S*. Typhimurium D23580ΔΦ (for P22 phage enumeration); phage-free *S. aureus* RN451 (for phi80α and phi11 phage enumeration) were back-diluted 1:100 into LB (for *E. coli* and *S*. Typhimurium) or BHI (for *S. aureus*) and incubated at 37 °C. At OD_600_ 0.3-0.5*, E. coli* and *S*. Typhimurium cultures were diluted 1:10 into molten LB-agar (0.6%) supplemented with 10 mM MgSO_4_ and 0.2% maltose and poured onto a LB-agar (1.5%) plate. For *S. aureus*, cultures were back-diluted 1:10 into molten tryptic soy agar (0.6%) supplemented with 10mM CaCl_2_ and poured onto a denser layer (1.5%) of the same agar.

### Bioinformatic analyses

NCBI tBLASTn (nr/nt database, expect threshold = 0.05, word size = 6, BLOSUM62 matrix) was used to identify *clbS* genes that match *E. coli* ClbS (WP_000290498) but that are found outside of *pks* clusters. The more distantly related ClbS-like proteins examined in this study (Figure 3d) were compiled from BLASTp results using *E. coli* ClbS as the query. After excluding entries that occur in genomes with *pks* clusters, the isolation source of the remaining hits was considered in identifying bee gut and human-associated isolates. Other members in the representative panel selected for cloning and heterologous expression were chosen heuristically and to cover the range in percent identities returned by the BLAST search (spanning *M. theicola* having 80% pairwise identity and *B. longum* with 26.8% pairwise identity to *E. coli* ClbS). The genomes encoding *clbS*-like genes in the representative panel were submitted to PHASTER^41^ for identification of prophage regions. Genes encoded by predicted intact prophages (score >90) were further analyzed by domain analysis (InterPro) for features matching the lambda repressor (DNA binding and peptidase domains), as mentioned in the main text and depicted in Extended Data Figure 6c.

### Quantification and statistical analysis

Data are presented as the mean ± std unless otherwise indicated in the figure legends. The number of independent biological replicates for each experiment is indicated for each experiment and included in the legend.

## Data and software availability

Unprocessed images and associated data generated in the course of this study are deposited on Zenodo (doi: 10.5281/zenodo.4683078). Other experimental data that support the findings of this study are available by reasonable request from the corresponding author.

## Acknowledgments

We thank all members of the Balskus group for insightful discussion. We thank Kai Papenfort (FSU-Jena) for helpful feedback and the Penadés lab (Imperial College London) for sharing *S. aureus* strains. This work was supported by the National Institutes of Health Grant R01 CA208834. JWHW was supported by the A*STAR NSS (PhD) predoctoral fellowship. SVO and MB were partially supported by the NIH NIGMS award R35GM133700, the David and Lucile Packard Foundation, and the Pew Charitable Trusts. The content is solely the responsibility of the authors and does not necessarily represent the official views of the funders.

## Author contributions

JES, JWHW, and EPB conceived of the project. JES and JWHW constructed strains and performed all experiments. JES, JWHW, SVO, MB, and EPB interpreted data and provided critical feedback. JES, JWHW, and EPB wrote the paper.

## Additional information

Supplementary Information is available for this paper. Correspondence and requests for materials should be addressed to.

## Extended Data

**Extended Data Figure 1.**
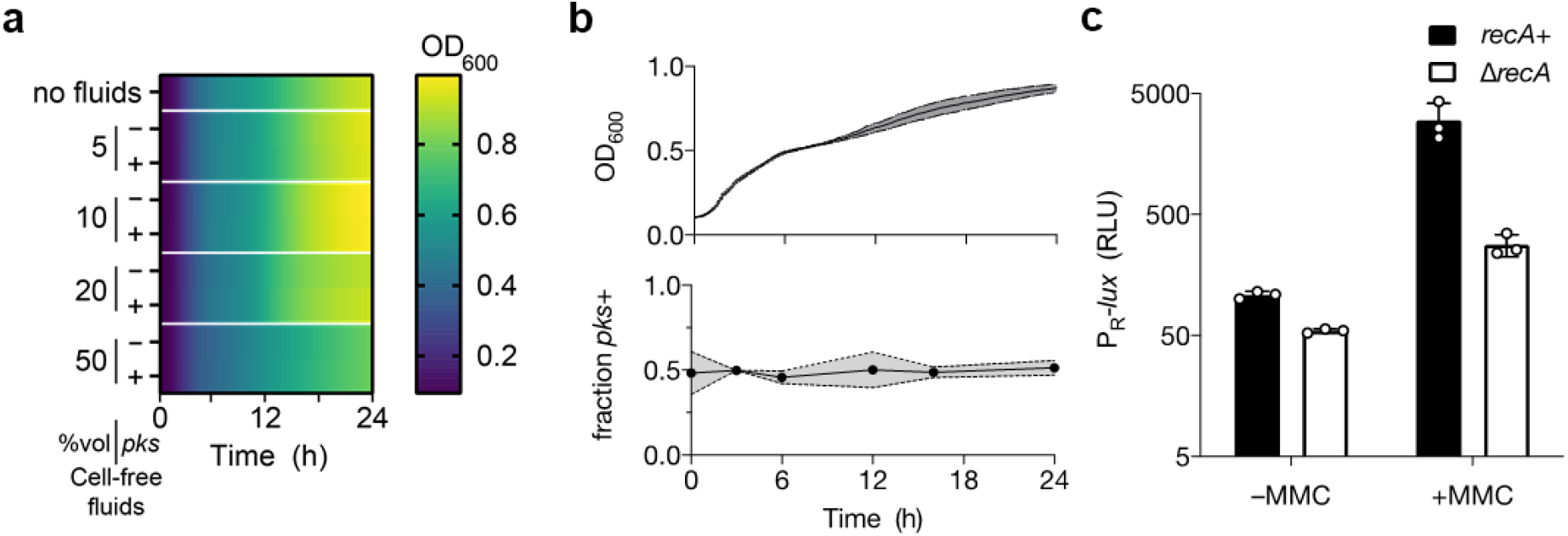
Colibactin production does not generally inhibit bacterial growth but induces DNA damage. **a** Growth of *pks*^−^ *E. coli* grown in the presence of the indicated relative volume of cell-free fluids from overnight cultures of *pks*^+^ *E. coli, pks*^−^ *E. coli* or without cell-free fluids added (top row). **b** Growth and frequency of *pks*^−^ and *pks*^+^ *E. coli* in co-culture as in Figure 1a but with the *pks/lacZ* combination swapped. Upper, total culture density of *pks^−^/lacZ^+^ E. coli* co-cultured with *pks*^+^/*lacZ*^−^ *E. coli* at a starting ratio of 1:1; lower, the proportion of *lacZ*^+^ within the same co-culture based on differential blue-white plating over time. **c** P_R_-*lux* output in *recA*^+^ (black) and Δ*recA* (white) *E. coli* harboring the reporter plasmid in the absence and presence of 100 ng mL^−1^ MMC. For c, RLU as in 1e. Data represented as mean of n = 3 biological replicates (a) or as mean ± SD with n = 3 biological replicates (b and c).

**Extended Data Figure 2.**
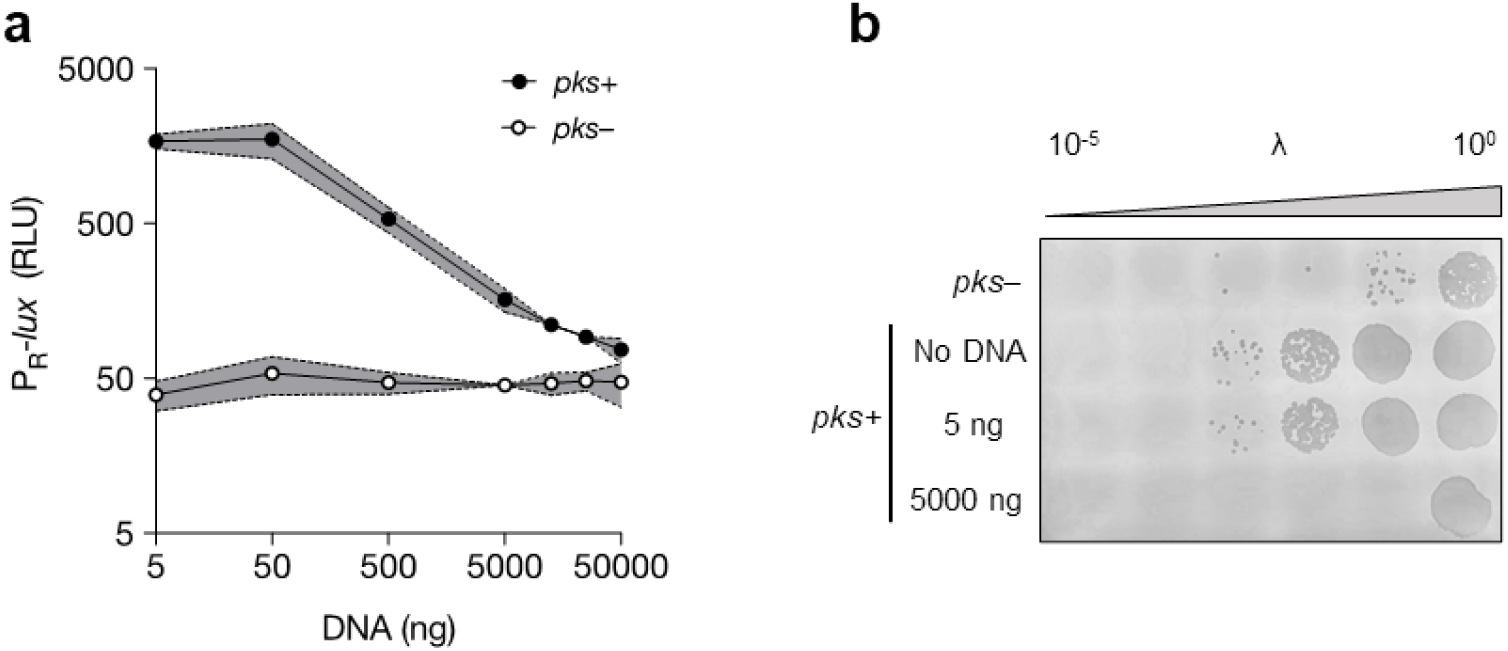
Addition of extracellular DNA inhibits phage induction caused by colibactin. **a** P_R_-*lux* output from reporter cells co-cultured with either *pks*^+^ or *pks*^−^ *E. coli* and the indicated amount of herring sperm DNA. **b** Plaque assays of the analogous incubations as in (a) but with a lambda lysogen used in place of the reporter strain. For a, RLU as in 1e. Data represented as mean ± SD with n = 3 biological replicates (a) or n = 3 biological replicates from which a single representative image is shown (b).

**Extended Data Figure 3.**
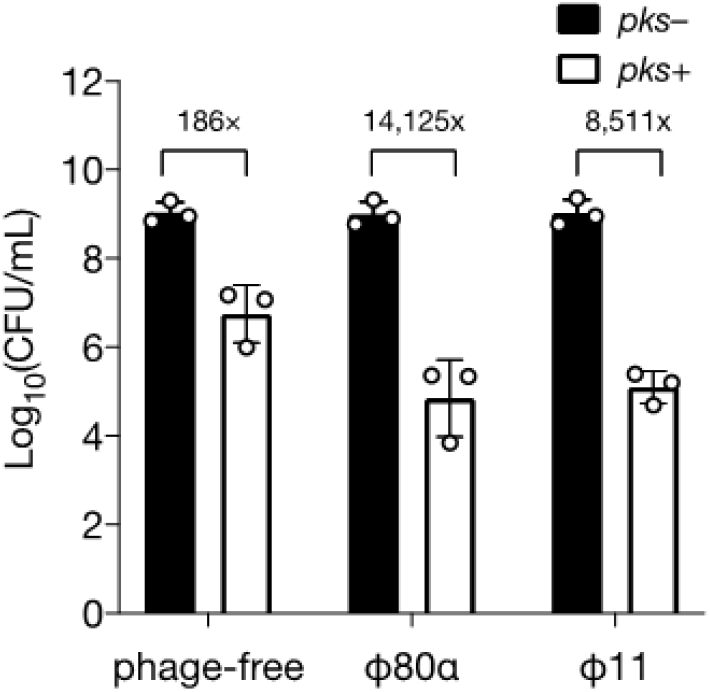
*S. aureus* harboring prophages are more susceptible to colibactin. Colony forming units of lysogenic and non-lysogenic (phage-free) *S. aureus* after being co-cultured with *pks*^+^ and *pks*^−^ *E. coli*. Data represented as mean ± SD with n = 3 biological replicates.

**Extended Data Figure 4.**
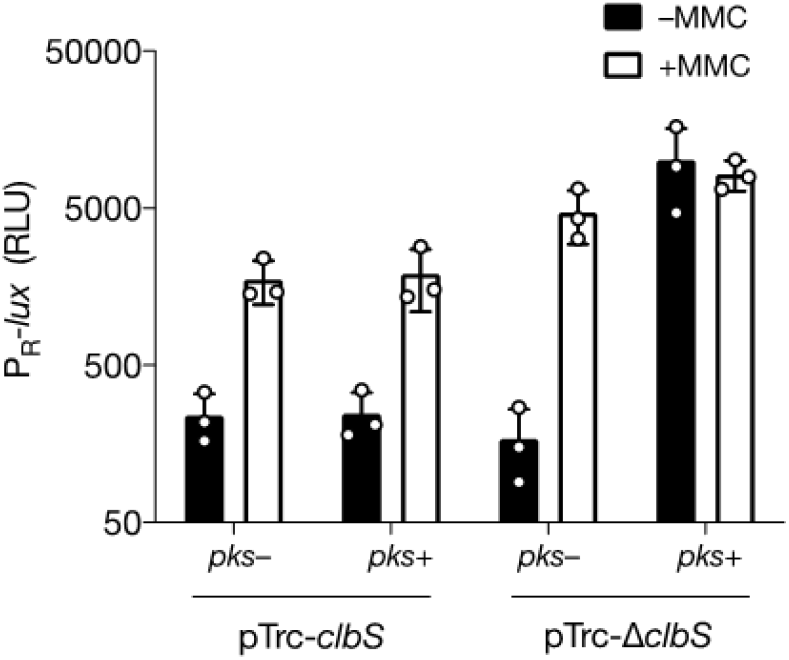
ClbS provides colibactin-specific protection from DNA damage. P_R_-*lux* reporter output in in the absence and presence of 100 ng mL^−1^ MMC in *E. coli* harboring the P_R_-*lux* reporter plasmid, the indicated second plasmid (*pTrc-clbS* or pTrc-Δ*clbS*), and co-cultured with *pks*^−^ or *pks*^+^ *E. coli*. RLU as in 1e. Data represented as mean ± SD with n = 3 biological replicates.

**Extended Data Figure 5.**
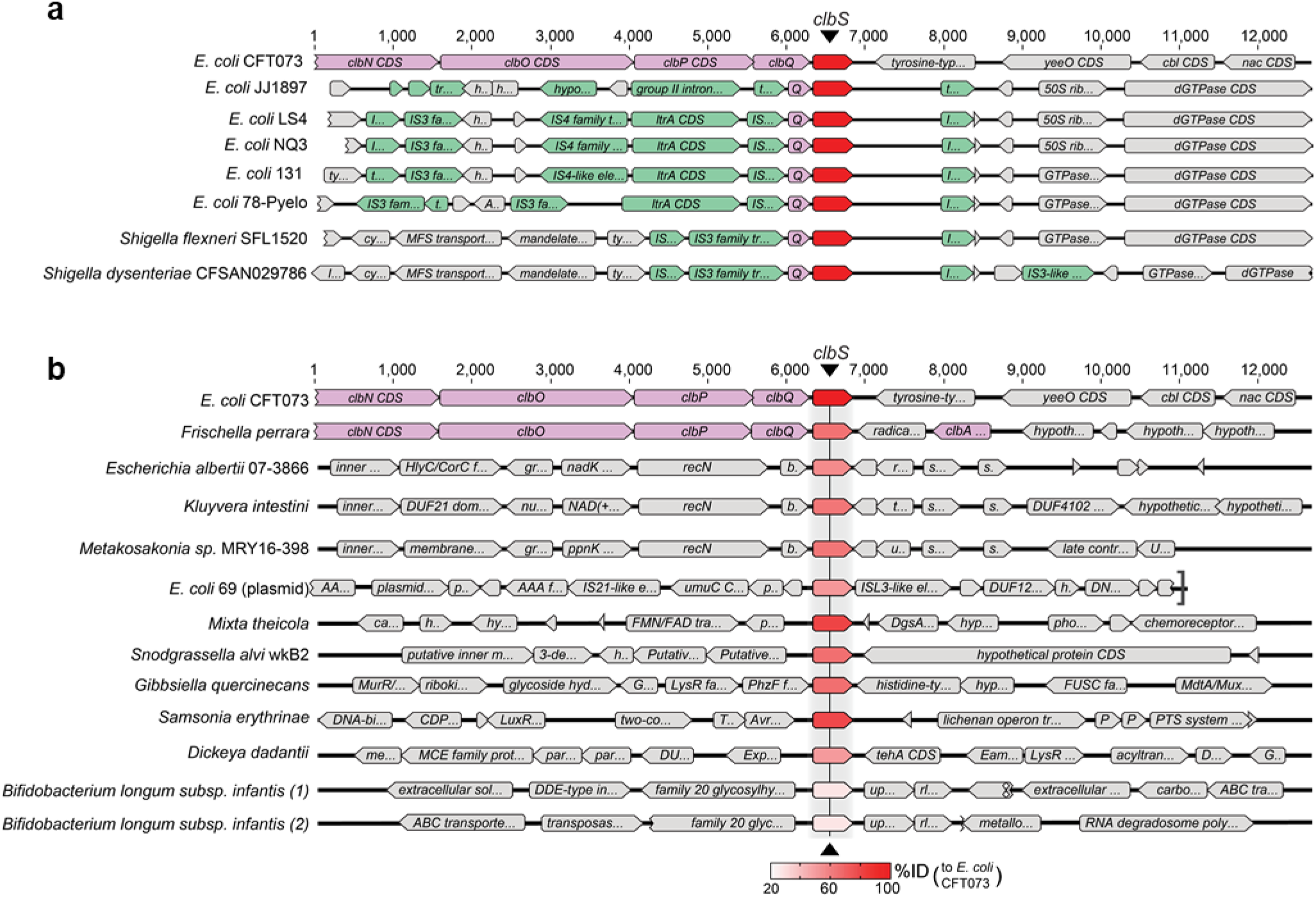
*clb*S-like genes are present in genomes of diverse bacteria, including those lacking *pks*-biosynthetic genes. **a** Genomic context of *clbS* found within the *E. coli pks* cluster encoded by a known colibactin producing isolate (CFT073) as compared to *pks*^−^ isolates of *E. coli* that lack the colibactin biosynthetic genes but contain an identical *clbS* coding sequence and truncated *clbQ* in regions flanked with predicted transposase-associated genes (green-colored genes). **b** Genomic organization surrounding *clbS*-like genes encoded by diverse bacteria identified in this study. Purple-colored genes denote the known *pks* biosynthetic genes. *E. coli* CFT073 and *F. perrara* were previously known to carry *pks*-associated *clbS*. In both panels, red-colored genes denote *clbS*. The saturation of red for each *clbS* is proportional to the percent identity in amino acid sequence of that gene relative to *E. coli* CFT073, as indicated in the key. Numbering at top in each panel refers to sequence length in base pairs.

**Extended Data Figure 6.**
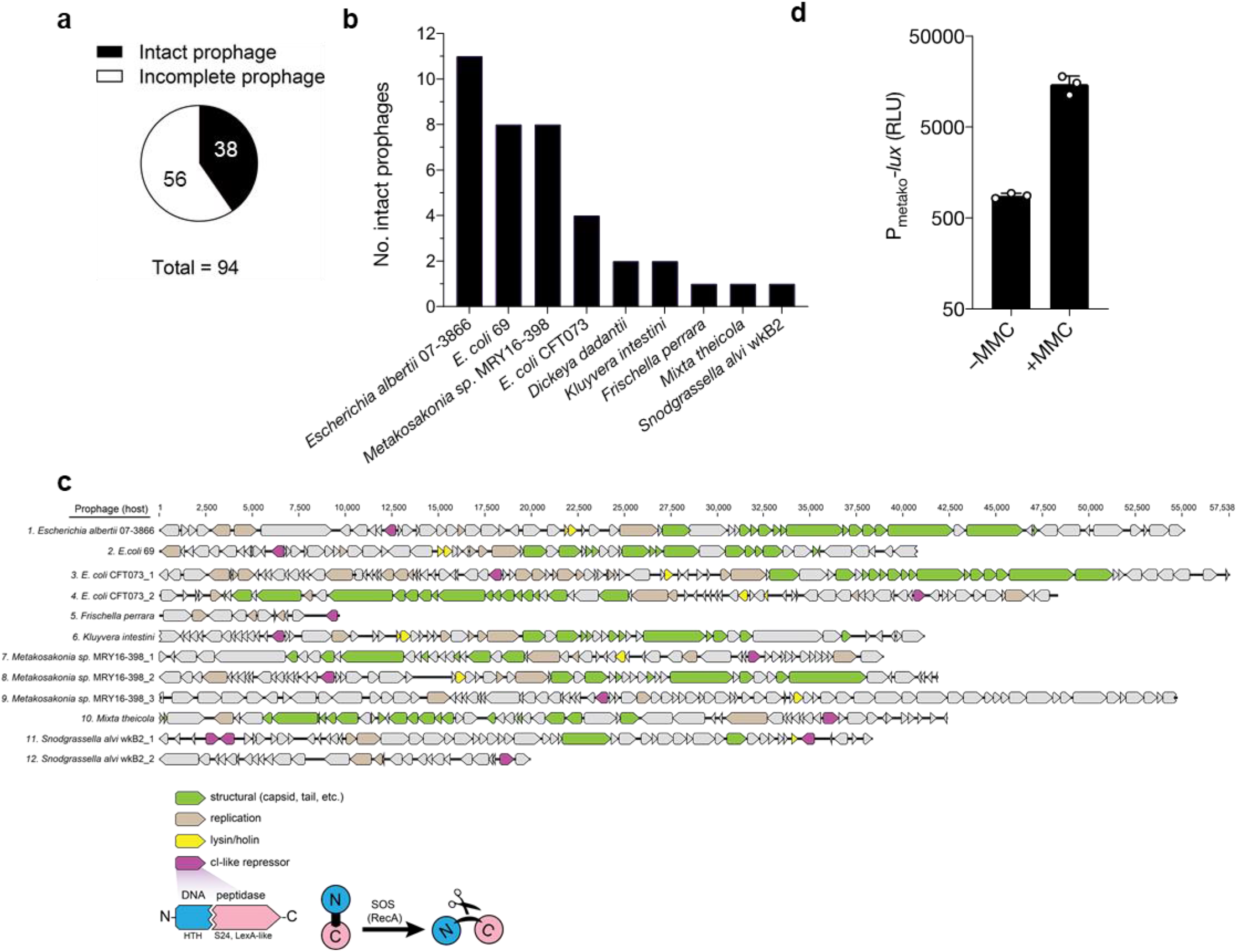
Prophages with predicted DNA-damage responsive repressors co-occur in *clbS*-encoding bacteria. **a** Distribution of PHASTER-predicted prophage regions present in the 12 bacterial genomes that encode the *clbS-like* genes tested in Figure 3c (genomic context for each shown in Extended Data Figure 5b). A total of 94 prophage regions were predicted, 38 of which are considered to be intact phages. **b** Number and distribution of intact prophages within each bacterial species from (a). **c** Organization of predicted intact phage genomes that encode prototypical DNA-damage responsive repressors (12 from the 38 intact phages identified in a and b). Genes colored according to predicted function, designated in the key. Numbering above genomes denotes prophage genome size in base pairs. Domain analysis was used to predict the cI-like repressor (maroon genes) on the basis that it harbors a helix-turn-helix DNA binding domain (blue, N-terminal domain) and a LexA-like, S24 peptidase domain (pink, C-terminal domain). The same two-domain architecture is found in the lambda cl repressor and confers an autoproteolytic mechanism in which the repressor is cleaved in the presence of a DNA-damage-induced, RecA-active protein complex, leading to phage lysis. **d** P_Metako_-*lux* output in *recA*^+^ *E. coli* harboring the reporter plasmid in the absence and presence of 100 ng mL^−1^ MMC. RLU as in 1e. Data represented as mean of n = 3 biological replicates.

## References

1. Nougayrède, J.-P. et al. Escherichia coli Induces DNA Double-Strand Breaks in Eukaryotic Cells. Science 313, 848–851 (2006).

2. Auvray, F. et al. Insights into the acquisition of the pks island and production of colibactin in the Escherichia coli population. Microb. Genomics 7, 000579.

3. Dougherty, M. W. & Jobin, C. Shining a Light on Colibactin Biology. Toxins 13, 346 (2021).

4. Newman, D. J. & Cragg, G. M. Natural Products as Sources of New Drugs over the Nearly Four Decades from 01/1981 to 09/2019. J. Nat. Prod. 83, 770–803 (2020).

5. Meksuriyen, D. & Cordell, G. A. Biosynthesis of Staurosporine, 1. 1 H- and 13 C-nmr Assignments. J. Nat. Prod. 51, 884–892 (1988).

6. Pishchany, G. & Kolter, R. On the possible ecological roles of antimicrobials. Mol. Microbiol. 113, 580–587 (2020).

7. Sengupta, S., Chattopadhyay, M. K. & Grossart, H.-P. The multifaceted roles of antibiotics and antibiotic resistance in nature. Front. Microbiol. 4, (2013).

8. Bondarev, V. et al. The genus Pseudovibrio contains metabolically versatile bacteria adapted for symbiosis. Environ. Microbiol. 15, 2095–2113 (2013).

9. Engel, P., Vizcaino, M. I. & Crawford, J. M. Gut Symbionts from Distinct Hosts Exhibit Genotoxic Activity via Divergent Colibactin Biosynthesis Pathways. Appl. Environ. Microbiol. 81, 1502–1512 (2015).

10. Moretti, C. et al. Erwinia oleae sp. nov., isolated from olive knots caused by Pseudomonas savastanoi pv. savastanoi. Int. J. Syst. Evol. Microbiol. 61, 2745–2752 (2011).

11. Bossuet-Greif, N. et al. The Colibactin Genotoxin Generates DNA Interstrand Cross-Links in Infected Cells. mBio 9, (2018).

12. Xue, M., Wernke, K. M. & Herzon, S. B. Depurination of Colibactin-Derived Interstrand Cross-Links. Biochemistry 59, 892–900 (2020).

13. Arthur, J. C. et al. Intestinal Inflammation Targets Cancer-Inducing Activity of the Microbiota. Science 338, 120–123 (2012).

14. Wilson, M. R. et al. The human gut bacterial genotoxin colibactin alkylates DNA. Science 363, (2019).

15. Dziubańska-Kusibab, P. J. et al. Colibactin DNA-damage signature indicates mutational impact in colorectal cancer. Nat. Med. 26, 1063–1069 (2020).

16. Pleguezuelos-Manzano, C. et al. Mutational signature in colorectal cancer caused by genotoxic pks + E. coli. Nature 580, 269–273 (2020).

17. Faïs, T., Delmas, J., Barnich, N., Bonnet, R. & Dalmasso, G. Colibactin: More Than a New Bacterial Toxin. Toxins 10, (2018).

18. Williams, P. C., Wernke, K. M., Tirla, A. & Herzon, S. B. Employing chemical synthesis to study the structure and function of colibactin, a “dark matter” metabolite. Nat. Prod. Rep. 37, 1532–1548 (2020).

19. Balskus, E. P. Colibactin: understanding an elusive gut bacterial genotoxin. Nat. Prod. Rep. 32, 1534–1540 (2015).

20. Jiang, Y. et al. Reactivity of an Unusual Amidase May Explain Colibactin’s DNA Cross-Linking Activity. J. Am. Chem. Soc. 141, 11489–11496 (2019).

21. Xue, M. et al. Structure elucidation of colibactin and its DNA cross-links. Science 365, (2019).

22. Tronnet, S. et al. The Genotoxin Colibactin Shapes Gut Microbiota in Mice. mSphere 5, (2020).

23. Faïs, T. et al. Antibiotic Activity of Escherichia coli against Multiresistant Staphylococcus aureus. Antimicrob. Agents Chemother. 60, 6986–6988 (2016).

24. Secher, T., Samba-Louaka, A., Oswald, E. & Nougayrède, J.-P. Escherichia coli Producing Colibactin Triggers Premature and Transmissible Senescence in Mammalian Cells. PLOS ONE 8, e77157 (2013).

25. Bossuet-Greif, N. et al. Escherichia coli ClbS is a colibactin resistance protein. Mol. Microbiol. 99, 897–908 (2016).

26. Tripathi, P. et al. ClbS Is a Cyclopropane Hydrolase That Confers Colibactin Resistance. J. Am. Chem. Soc. 139, 17719–17722 (2017).

27. Dubois, D. et al. ClbP is a prototype of a peptidase subgroup involved in biosynthesis of nonribosomal peptides. J. Biol. Chem. 286, 35562–35570 (2011).

28. Brotherton, C. A. & Balskus, E. P. A Prodrug Resistance Mechanism Is Involved in Colibactin Biosynthesis and Cytotoxicity. J. Am. Chem. Soc. 135, 3359–3362 (2013).

29. Ofir, G. & Sorek, R. Contemporary Phage Biology: From Classic Models to New Insights. Cell 172, 1260–1270 (2018).

30. Gimble, F. S. & Sauer, R. T. λ Repressor mutants that are better substrates for RecA-mediated cleavage. J. Mol. Biol. 206, 29–39 (1989).

31. Vizcaino, M. I. & Crawford, J. M. The colibactin warhead crosslinks DNA. Nat. Chem. 7, 411–417 (2015).

32. Huo, L. et al. Heterologous expression of bacterial natural product biosynthetic pathways. Nat. Prod. Rep. 36, 1412–1436 (2019).

33. Yang, Y., Gharaibeh, R. Z., Newsome, R. C. & Jobin, C. Amending microbiota by targeting intestinal inflammation with TNF blockade attenuates development of colorectal cancer. Nat. Cancer 1, 723–734 (2020).

34. Sheldon, P. J., Mao, Y., He, M. & Sherman, D. H. Mitomycin resistance in Streptomyces lavendulae includes a novel drug-binding-protein-dependent export system. J. Bacteriol. 181, 2507–2512 (1999).

35. Martin, T. W. et al. Molecular Basis of Mitomycin C Resistance in Streptomyces: Structure and Function of the MRD Protein. Structure 10, 933–942 (2002).

36. Tripathi, P. & Bruner, S. D. Structural Basis for the Interactions of the Colibactin Resistance Gene Product ClbS with DNA. Biochemistry (2021) doi:10.1021/acs.biochem.1c00201.

37. Putze, J. et al. Genetic Structure and Distribution of the Colibactin Genomic Island among Members of the Family Enterobacteriaceae. Infect. Immun. 77, 4696–4703 (2009).

38. Sekizuka, T. et al. Complete Genome Sequence of blaIMP–6-Positive Metakosakonia sp. MRY16-398 Isolate From the Ascites of a Diverticulitis Patient. Front. Microbiol. 9, (2018).

39. Tetz, G. et al. Complete Genome Sequence of Kluyvera intestini sp. nov., Isolated from the Stomach of a Patient with Gastric Cancer. Genome Announc. 5, (2017).

40. Kwong, W. K. & Moran, N. A. Gut microbial communities of social bees. Nat. Rev. Microbiol. 14, 374–384 (2016).

41. Arndt, D. et al. PHASTER: a better, faster version of the PHAST phage search tool. Nucleic Acids Res. 44, W16–W21 (2016).

42. Martin, P. et al. Interplay between Siderophores and Colibactin Genotoxin Biosynthetic Pathways in Escherichia coli. PLOS Pathog. 9, e1003437 (2013).

43. Massip, C. et al. Deciphering the interplay between the genotoxic and probiotic activities of Escherichia coli Nissle 1917. PLOS Pathog. 15, e1008029 (2019).

44. Lindsay, J. A. Genomic variation and evolution of Staphylococcus aureus. Int. J. Med. Microbiol. 300, 98–103 (2010).

45. Xia, G. & Wolz, C. Phages of Staphylococcus aureus and their impact on host evolution. Infect. Genet. Evol. 21, 593–601 (2014).

46. Zhang, X. et al. Quinolone Antibiotics Induce Shiga Toxin-Encoding Bacteriophages, Toxin Production, and Death in Mice. J. Infect. Dis. 181, 664–670 (2000).

47. Hendrix, R. W., Smith, M. C. M., Burns, R. N., Ford, M. E. & Hatfull, G. F. Evolutionary relationships among diverse bacteriophages and prophages: All the world’s a phage. Proc. Natl. Acad. Sci. 96, 2192–2197 (1999).

48. Mushegian, A. R. Are There 1031 Virus Particles on Earth, or More, or Fewer? J. Bacteriol. 202, (2020).

49. Duerkop, B. A., Clements, C. V., Rollins, D., Rodrigues, J. L. M. & Hooper, L. V. A composite bacteriophage alters colonization by an intestinal commensal bacterium. Proc. Natl. Acad. Sci. 109, 17621–17626 (2012).

50. Oh, J.-H. et al. Dietary Fructose and Microbiota-Derived Short-Chain Fatty Acids Promote Bacteriophage Production in the Gut Symbiont Lactobacillus reuteri. Cell Host Microbe 25, 273–284.e6 (2019).

51. Cummings, J. H., Pomare, E. W., Branch, W. J., Naylor, C. P. & Macfarlane, G. T. Short chain fatty acids in human large intestine, portal, hepatic and venous blood. Gut 28, 1221–1227 (1987).

52. Ferraris, R. P., Choe, J. & Patel, C. R. Intestinal Absorption of Fructose. Annu. Rev. Nutr. 38, 41–67 (2018).

53. Pavlova, S. I., Kiliç, A. O., Mou, S. M. & Tao, L. Phage Infection in Vaginal Lactobacilli: An In Vitro Study. Infectious Diseases in Obstetrics and Gynecology https://www.hindawi.com/journals/idog/1997/793172/abs/ (1997) doi:10.1155/S1064744997000094.

54. Selva, L. et al. Killing niche competitors by remote-control bacteriophage induction. Proc. Natl. Acad. Sci. 106, 1234–1238 (2009).

55. Kronheim, S. et al. A chemical defence against phage infection. Nature 564, 283–286 (2018).

56. Jiang, Z., Wei, J., Liang, Y., Peng, N. & Li, Y. Aminoglycoside Antibiotics Inhibit Mycobacteriophage Infection. Antibiotics 9, (2020).

57. Kever, L. et al. Aminoglycoside antibiotics inhibit phage infection by blocking an early step of the phage infection cycle. bioRxiv 2021.05.02.442312 (2021) doi:10.1101/2021.05.02.442312.

58. Jancheva, M. & Böttcher, T. A Metabolite of Pseudomonas Triggers Prophage-Selective Lysogenic to Lytic Conversion in Staphylococcus aureus. J. Am. Chem. Soc. (2021) doi:10.1021/jacs.1c01275.

59. Silpe, J. E. & Bassler, B. L. A Host-Produced Quorum-Sensing Autoinducer Controls a Phage Lysis-Lysogeny Decision. Cell 176, 268–280.e13 (2019).

60. Gregory, A. C., Zablocki, O., Howell, A., Bolduc, B. & Sullivan, M. B. The human gut virome database. (2019) doi:10.1101/655910.

61. Ma, Y., You, X., Mai, G., Tokuyasu, T. & Liu, C. A human gut phage catalog correlates the gut phageome with type 2 diabetes. Microbiome 6, 24 (2018).

62. Tetz, G., Brown, S. M., Hao, Y. & Tetz, V. Type 1 Diabetes: an Association Between Autoimmunity, the Dynamics of Gut Amyloid-producing E. coli and Their Phages. Sci. Rep. 9, 9685 (2019).

63. Zhao, G. et al. Intestinal virome changes precede autoimmunity in type I diabetes-susceptible children. Proc. Natl. Acad. Sci. 114, E6166–E6175 (2017).

64. Han, M., Yang, P., Zhong, C. & Ning, K. The Human Gut Virome in Hypertension. Front. Microbiol. 9, (2018).

65. Norman, J. M. et al. Disease-Specific Alterations in the Enteric Virome in Inflammatory Bowel Disease. Cell 160, 447–460 (2015).

66. Hannigan, G. D., Duhaime, M. B., Ruffin, M. T., Koumpouras, C. C. & Schloss, P. D. Diagnostic Potential and Interactive Dynamics of the Colorectal Cancer Virome. mBio 9, (2018).

67. Clooney, A. G. et al. Whole-Virome Analysis Sheds Light on Viral Dark Matter in Inflammatory Bowel Disease. Cell Host Microbe 26, 764–778.e5 (2019).

