## Supplementary material for "The gut bacterial natural product colibactin triggers induction of latent viruses in diverse bacteria": Table S1

### SUPPORTING INFORMATION

**Supplementary Table 1.** tBLASTn result of top 230 *clbS*-like genes encoding proteins matching *E. coli* ClbS, WP_000290498 (available as a separate excel).

**Supplementary Table 2.** Strains used in this study.

**Supplementary Table 3.** Plasmids used in this study.

**Supplementary Table 4.** Gene blocks and oligonucleotides used in this study.

**Supplementary Table 1|** tBLASTn result of top 250 *clbS* genes encoding proteins matching *E. coli* ClbS, WP_000290498 (available as a separate excel).

**Supplementary Table 2|** Strains used in this study.

| Strain | Genotype | Reference |
|---|---|---|
| *E. coli* DH10B | *F– mcrA Δ(mrr-hsdRMS-mcrBC) φ80lacZΔM15 ΔlacX74 recA1 araD139 Δ(ara-leu)7697 galU galK λ– rpsL(Str$^R$) endA1 nupG* | NEB |
| *E. coli* BW25113 | *lacIq, rrnBT14, ΔlacZWJ16, hsdR514,ΔaraBADAH33, ΔrhaBADLD78* | Baba et al. 2006 |
| *E. coli* BW25113 λ | BW25113 lysogenic for λ | This study |
| *E. coli* BW25113 *lamB::kan* | *lacIq, rrnBT14, ΔlacZWJ16, hsdR514,ΔaraBADAH33, ΔrhaBADLD7 ΔlamB732::kan* | Baba et al. 2006 (JW3996-1) |
| *E. coli* MG1655 KIlac | MG1655 with Lac Operon removed from native locus and re-integrated at the Tn7 locus | Eames et al. 2012 (Addgene: #52696) |
| *E. coli* MG1655 (delta-Z) | KIlac with *lacZ* deletion | Eames et al. 2012 (Addgene: #52706) |
| *E coli* NC101 | Murine isolate of *E. coli* (causes colitis in gnotobiotic IL-10−/− mice) | Tomkovich et al. 2017 |
| *E. coli* NC101 Δ*clbP* | Δ*clbP* mutant of *E. coli* NC101 | Tomkovich et al. 2017 |
| *S.* Typhimurium D23580 ΔΦ (JH3949) | D23580, cured of 5 native prophages | Owen et al. 2017 |
| *S.* Typhimurium D23580 ΔΦ [P22] (SSO128) | D23580 ΔΦ lysogenic for P22 | Owen et al. 2021 |
| *S. aureus* RN450 | NCTC8325 cured of Φ11, Φ12, Φ13 | Novick R 1967 |
| *S. aureus* RN10359 | RN450 lysogenic for Φ80a | Ubeda et al. 2007 |
| *S. aureus* RN451 | RN450 lysogenic for Φ11 | Novick 1967 |

**Supplementary Table 3 |** Plasmids used in this study.

| Plasmid | Relevant fragment | Resistance | Reference |
|---|---|---|---|
| JSS-2347 | empty BAC (pBeloBAC11) | Cm | NEB |
| JSS-2348 | BAC-*pks* | Cm | Gift of Bonnet Lab |
| JSS-2367 | BAC-*pks* Δ*clbP* | Kan Cm | Gift of Bonnet Lab |
| JSS-2435 | empty BAC2 (cos sites removed) | Cm | This study |
| JSS-2403 | $P_R$-*lux* | Kan | This study |
| JSS-2587 | pBR322-empty (*ampR*) | Amp | This study |
| JSS-2436 | pEVS143-empty (*kanR*) | Kan | This study |
| JSS-2442 | pEVS143-empty (*kanR-cmR*) | Kan Cm | This study |
| JSS-2470 | pTrc-*clbS* | Amp | This study |
| JSS-2482 | pTrc-Δ*clbS* (pTrc-*clbS* with *clbS* deleted) | Amp | This study |
| JSS-2522 | pTrc-$clbS_{Dickeya}$ (*Dickeya dadantii*) | Amp | This study |
| JSS-2523 | pTrc-$clbS_{Mixta}$ (*Mixta theicola*) | Amp | This study |
| JSS-2528 | pTrc-$clbS_{Ecoli69}$ (*Escherichia coli 69,* plasmid) | Amp | This study |
| JSS-2529 | pTrc-$clbS_{Sond}$ *(Snodgrassella alvi wkB2)* | Amp | This study |
| JSS-2530 | pTrc-$clbS_{Gibsiella}$ (*Gibbsiella quercinecans* DSM 25889) | Amp | This study |
| JSS-2531 | pTrc-$clbS_{Samsonia}$ (*Samsonia erythrinae* DSM 16730) | Amp | This study |
| JSS-2532 | pTrc-$clbS_{Frischella}$ (*Frischella perrara*) | Amp | This study |
| JSS-2555 | pTrc-$clbS_{Bifido}$ (*Bifidobacterium longum subsp. infantis*) | Amp | This study |
| JSS-2556 | pTrc-$clbS_{Ealbertii}$ (*Escherichia albertii* CB9786) | Amp | This study |
| JSS-2557 | pTrc-$clbS_{Metako}$ (*Metakosakonia sp.* MRY16-398) | Amp | This study |
| JSS-2576 | $P_{Metako}$-*lux* | Kan | This study |

**Supplementary Table 4 |** Gene blocks and oligonucleotides used in this study.

| ID | Sequence (5-3’) |
|---|---|
| JSgblock-0133 (*Dickeya dadantii*) | ATATTAATGTATCGATTAAATAAGGAGGAATAAACCATGGGCGTGCCACAAACAAAATCCGAACTACT<br>TTTCGCTATTGATAAAAATTTTACGAAATTAATCGGTTACCTCAATGCGATTCCGCCAGAGCTCACTG<br>CGGATAAATCGCTGGATGGCCATGCAAAAGGCACGGAGATGAGTGTTTGTGAGCTTGTTTCGTATTT<br>GCTGGGATGGAACACTCTGGTTGTAAAATGGATTAGCGGTGACGCAACAGGGCAACCTGTTGATTT<br>CCCGGAAACCGGTTACAAGTGGAATCAGCTCGGGCTTCTGGCGCAGAAATTTTACTTTGATTACAAA<br>GAATTAAATTACCAGACGTTAATCAATAACCTTCAGTCGGTAAAAGACGAAATAGTGAGGCTGATTGA<br>TGAGCGCACTGATGAAGATCTGTATGGCAAGCCATGGTATGGCAAATGGACAATGGGACGTATGAT<br>CTCATTGAACACGTCATCACCCTATGCCAATGCTAATGGAAGATTAAGGCAGTGGGCAAAGAACAAT<br>CACCTGCGTTTAAAATGAGTTTTGGCGGATGAGAGAAGATTTTCAGCCTGATAC |
| JSgblock-0134 (*Frischella perrara*) | ATATTAATGTATCGATTAAATAAGGAGGAATAAACCATGGCAGTACCTGAATCTAAAACTGAGTTAAT<br>AGAAGCGATTAATAAAAATTATGCTTCGTTAGTCAAAAAGCTATCAGCTGTACCAGCAGAAAAGGCTT<br>TCCAACCATTAATGGAGGGGCATGCCAAAGGCACGACTATGAGTGCAGCACAACTTGTTTCTTATTT<br>AATAGGTTGGGGCGAATTAGTCTTATCTTGGCATGCACAAGAAGCACATGGGCATAGCATCGTTTTT<br>CCTGAAGTGGGCTTTAAATGGAATGAATTAGGAAAACTAGCACAGAAATTTTATCAAGACTATCAAAA<br>CATCACTGATTACAAGGAGCTATTACAACGGTTAGAAAAAAATAAAAATGATCTTATTAAATTGATTGA<br>AGGTTTTACGAATAATGATTTATATGGTTCGAGTTGGTGTGAAAAATACACTCGAGGACGAATGATTC<br>AGCTAAATACGTCATCACCGTATAGAAATGCAAGTGGTCGCATCGCGACATTCCTCAAAACTATCGG<br>CAATTAAGTTTTGGCGGATGAGAGAAGATTTTCAGCCTGATAC |
| JSgblock-0135 (*Mixta theicola*) | ATATTAATGTATCGATTAAATAAGGAGGAATAAACCATGGCGGTACCGGGATCAAAAGCTGAATTAAT<br>TAAAGCGATCAATAGTAATTTCGCGTTGCTGAATAAAAAGCTGGAGGCTATTGCGCCAGCGCTTGCC<br>TTTGAGCCGCTTATGGAAGGGCATGCCAAAGGGAGCACGATGAGCGTGGCGCAGCTGGTCTCCTG<br>GCTGATTGGCTGGGGCGAATTGGTGTTGCACTGGCATAACCAAGAGGCGAAAGGGGAAACCATCGT<br>GTTTCCTGCAGAGGGCTATAAATGGAATGAGCTGGGACAGCTGGCGCAGAAATTCTACCGTGACTA<br>TGAAGACATCAACAATTATGAAACCTTGTTGGCGCGGTTAAGGGAGAACAAACAGCGGCTCTTGATG<br>CTGACTGAGCAATTTAGCAACGATGAACTTTACGGTACGCCCTGGTATGGCAAATGGACGCGAGGA<br>CGAATGATTCAATTTAATACCGCCTCGCCCTATAGAAATGCCGCAGGACGATTAAACAAATTGAAAAA<br>ACAAACTGCTGCGTAAGTTTTGGCGGATGAGAGAAGATTTTCAGCCTGATAC |
| JSgblock-0136 (*Escherichia coli* 69, plasmid) | ATATTAATGTATCGATTAAATAAGGAGGAATAAACCATGAGTGTGCCGCAAACAAAAGCTGAACTGCT<br>TTTAGCTATTGATAAAAATTTTAGTAAATTAATTAGTTACCTCAACACAATCCCACCAGAAATTACTTCA<br>GATAAATCAATGGACGGACACGCCAAAGGAACGGAGATGAGTGTTCGTGATCTCGTTTCGTATCTGC<br>TTGGATGGAATGCTCTTGTTGTAAAGTGGATCGCTTCTGATGCTAAAGGTCTGCCTGTCGATTTTCC<br>GGAAACTGGCTATAAATGGAATCAGCTTGGCCTTCTTGCTCAAAAATTTTACTCAGATTACAGTGAGT<br>TAAGTTATGAGTTGTTAGTAGCTGAACTTCAGACTGTAAAAAATGAGATTGTGAACCTTATTAATGATC<br>GTACCGATGATATTTTGTATGGAAGACCATGGTACACAAAATGGACGATGGGGAGAATGATCTCATT<br>TAACACATCTTCGCCTTACGCCAACGCTAATGGAAGATTAAGAAAGTGGGCAAAAAATAATAATATCA<br>GTTTAAAGTAAGTTTTGGCGGATGAGAGAAGATTTTCAGCCTGATAC |
| JSgblock-0137 (*Snodgrassella alvi* wkB2) | ATATTAATGTATCGATTAAATAAGGAGGAATAAACCATGGCAATACCAGAATCAAAACAGGAACTAAT<br>TGAAGCAATTAATAAAAACTATACACTACTGACCAAAAAGCTGGCTGCTGTACCTGAACAGAAAGCCT<br>ATCTGCCTCTGATGGAAGGCCATGCTAAAGGTACAATGATGAGTGCTGCACAGCTTGTGTCTTATTT<br>AATTGGCTGGGGAGAACTGGTGCTGTCATGGCATAAACAGGAACAGTCAGGGCAGAAAATTGCCTT<br>TCCGGAAGCAGGATATAAATGGAATCAACTGGGATTACTGGCACAGAAATTTTATCGTGATTATCAG<br>GACATTACCGATTTCAAACAGTTACTCTCGCTTCTGGAGACCAACAAAAAGAATCTGATTTCACTTAT<br>TGATAGCTTTAGTAATGAAGAGCTGTATGGCTCACCCTGGCATGAAAAATACACACGCGGACGCATG<br>ATTCAGTTTAATACCTCCTCCCCTTATAAAAATGCCACAGGACGATTGAATAAATTGTTGAAACAGAT<br>CAAAGAATAAGTTTTGGCGGATGAGAGAAGATTTTCAGCCTGATAC |
| JSgblock-0138 (*Gibbsiella quercinecans* DSM 25889) | ATATTAATGTATCGATTAAATAAGGAGGAATAAACCATGGCCGTGCCGGAATCAAAAGGCGCATTGA<br>TCAAAGCGGTGAACAGCAATTTCGCGCTGCTGATGAAAAAACTGGACGACATACCGGCAGAAAAGG<br>CGTTTGAGCCGCTGATGCCGGGGCATGCCAAGGGCAGCGTAATGAGCGTTGCCCAACTGGTGGCC<br>TATCTGATTGGCTGGGGGGAACTGGTTTTGTCCTGGCATGCGAAAGAAAGACAGGGCCTTGAAATC<br>GCCTTCCCCGAAGACGGTTTTAAGTGGAACCAATTGGGGTTGCTGGCGCAAAAGTTTTACCGTGATT<br>ATGCGCAGATTACGGACTACGGCGCCCTGCTGATGCGGCTTGCCGATAATAAACGGCAGATTATCG<br>CGTTGATTGACGGCTTTAGCGATGAGGCGCTCTACGGCCAGCCGTGGTACGGCAAATGGACGCGT<br>GGCCGGATGATTCAGTTCAACACGGCGTCACCGTATAAGAACGCCTGCGGCAGGCTAAACGCATTG<br>CGAAAAGTAGGCTGTGTCCCGTAAGTTTTGGCGGATGAGAGAAGATTTTCAGCCTGATAC |
| JSgblock-0139 (*Samsonia erythrinae* DSM 16730) | ATATTAATGTATCGATTAAATAAGGAGGAATAAACCATGGCGGTGCCAGAATCAAAAGATGAACTCAT<br>TAAAGCGATCAATAGTCACTTTGCGTTATTGCAGAAAAAGCTGGGGGCAGTCCTGCCTGAGCAGTCC<br>TTTGAACCGACAATGGAAGGGCATGCAAAAGGGAGCGTGATGAGCGTGGCCCAATTGGTGTCTTAT<br>CTTATTGGGTGGGGAGAATTGGTCTTGCACTGGCACGAACAAGAAGCGAAAGGGGAAACGATAAAC<br>TTTCCTGAAGAGGGCTATAAATGGAATGAATTGGGGCGCCTGGCGCAGAAGTTTTATCGCGACTATG<br>AAGACATAAAGGATTATGAAATCTTATTGTTGCGATTAAAGGAAAACAAACAGCAGCTTTTGACGCTG<br>ATTGAGCGCTTTAGTAACGAGGAACTTTACGGTAGTCCTTGGTATGGCAAATGGACGCGGGGGCGA<br>ATGATTCAATTTAATACCTCCTCGCCTTATAAAAATGCTTCAGGGCGATTAAATAAATTGAAAAAACAG<br>GCTCCGATATGAGTTTTGGCGGATGAGAGAAGATTTTCAGCCTGATAC |
| JSgblock-0140 (*Bifidobacterium longum subsp. infantis*) | ATATTAATGTATCGATTAAATAAGGAGGAATAAACC.ATGGCAGCCGGGAGGTGCATAGTGAGAACAT<br>ACGAGAACGCGCAGGAGCTCAAAAAGGAGATCGGCGCGGCGTTTCGAAAATACATTGCGGAGTTCG<br>ACGACATTCCGGAAGCCCTGAAAGACAAGCGCATCGACGAGGTCGAGAGGACCCCGGCGGAAAAC<br>CTCGCCTATCAGGTTGGTTGGACCACCCTGCTGCTGCAATGGGAGGATCGCGAGCGGAAGGGCCT<br>TCCGGTGCGAACGCCGTCGGACGAGTTCAAGTGGAACCAGCTCGGAAAGCTGTACCAGTGGTTCAC<br>CGACACCTACGCCCACCTTTCGCTGCGGGAGCTGAAGGGCATGTTGACCGACAACGTCGACGCCAT<br>ATACGCCATGATCGATGCGATGAGCGAGGACGAGCTGTTCAAGCCGCATATGAGGAAGTGGGCCG<br>ACGACGCCACCAAAACGGCGGTATGGGAAGTGTACAGGTTCATTCATGTGAACACCGTCGCCCCGT<br>TCGGATCGTTCAGGACGAAGATCCGCAAGTGGAAGCGGATGGCCCTATAG.GTTTTGGCGGATGAG<br>AGAAGATTTTCAGCCTGATAC |

| | |
|---|---|
| JSgblock-0142 (*Escherichia albertii* CB9786) | ATATTAATGTATCGATTAAATAAGGAGGAATAAACC.ATGTCTGTACCAGAAAGCAAAGCTGCACTGTT<br>GCTGGAAATGGAAAAATCATGGAATGGACTTACAAAAAAGTTATCTCGCATTCCAGAAGAAAAAGCTT<br>TTCTCATCACAATGGAAGGACATGCCAGCGGCACTGTGATGAGTGCGGCTAACCTGGTAAGCTATTT<br>AATAGGTTGGGGAGAGCAGGTTCTTACCTGGCACAGACAAGAAGAAGCAGGAATCCCCATTGATTTT<br>CCGGCTAAAGACCATAAGTGGAATGAACTTGGGAAACTAGCGCAGACATTTTACGCTAATTATTCCC<br>ATGTCACATCATGGTCCCAATTGTGCGAAATGTTAGAAACCAATCATCAGCAACTCAAATCGCTAGTG<br>GAACGCTACAGCGATAACGAGTTATACCATCATCCCTGGTATGGCAAATGGACGCGTGGTCGAATG<br>ATCCAATTTAATACCGTATCACCCTATAGAAATGCAAGCACACGTCTGAACGCCCTACTTAAACAATT<br>CTGAGTTTTGGCGGATGA.GAGAAGATTTTCAGCCTGATAC |
| JSgblock-0143 (*Metakosakonia* sp. MRY16-398 / *Kluyvera intestini* strain GT-16) | ATATTAATGTATCGATTAAATAAGGAGGAATAAACCATGGCCGTGCCAGAAAGCAAAACTGCCTTACT<br>GGAAGCAATGGAAAAGTCGGCAAGCGCCCTGCGCAAAAAACTCATCCGCATTCCGGCGGACATCGC<br>TTATCAGCAATGCCTGGAAGGGCACGTGGCTGGTAGCCGCATGAGTGTTGCTAATCTTGTCAGCTAT<br>CTGATTGGCTGGGGAGAGCAAGTTCTCTTCTGGCACCAACAGGAAGCCGCAGGCGAAGAAATAGAC<br>TTCCCGGCAAAAGGCTTTAAATGGAATGAACTGGGGAAGCTTGCGCAAAAATACTACGCTGATTACC<br>AGCATATTACGTCATGGCCAACGCTGCTTGCCATGCTGGAAGAGAATCAACAGCAACTGATATCACT<br>GGTGAATGGCTTTAGTGACGATGAGTTGTACCACCAGCTCTGGTACGGAAAATGGACGCGAGGACG<br>TATGATTCAATTCAACAGCGCCTCACCGTATAAGAATGCCAGCGCCAGGCTGAACAGTCTGTTAAAA<br>ACGCTCGCCGAAGCCTGACGCTCGCCGAAGCCTGAGTTTTGGCGGATGAGAGAAGATTTTCAGCCT<br>GATAC |
| JSgblock-0151 (*Metakosakonia* sp. MRY16-398_2 phage) | CGGGTACCGCTCGAGTTAATTAAGGCATAAAAAAACCAGCCGAAGCTGGTTTTTTATATGTATGAGT<br>CTCACCCCAAACGGTTGTATGACATTTGCCATCTACCGACAACTAAGCCCTGAATGTGAAACTCGCT<br>TTGATCAGTTTCAGATACAAACCATTTTTCGTAAGTTGAATTATCGCTGATGACTACCAGCTTGTCCTT<br>AAGCAGCTGTAAGCGTTTGATATGAAAGTTGTCACCATAAACGAACGCATAAATCCCATCACCTATCA<br>ACCGGTTTACCGTAACGTCTATAACGACAAGATCCCCAGGGAAAATAGTCCCGAGCATGCTATCGCC<br>GACAACCGTGCAGATTTTCAGGGAGGTAGGGCTACGCCCACCAAACATCCTTTTAGCTTCCTCTGGA<br>TCAAGTTCAATCGATTTAACTATTTCAGGATAGTCCATGTTCATCCTGCCACTTCCGCAACTGAACTC<br>AGTGTCCAGCACCTCAATTCGGTATTGTTGCTGATTTTTTGTCAGTGGATTTATGAAAAAGTTAGGCT<br>TGGTACCAGATGCATCAACATCTGCTCTCTTTGAATCCATCCAGCCATGGGGCAGGCCCATACTTGA<br>CTCAATTCTCCTAGCTATTATGTCACCAATGTTACGAGAGGCGTTAGAGCCAGATAACTGACTCAGTT<br>GAGAAGGGGGTATACCAATTTTTTCAGCAATTTGTGCCTTGGTATAACCTTCCACGATGTACTCACCA<br>ATCAACCTGTTTAGGTTCTGACGTCGAATGTTTTTTATGTCCATGACCAAATACTCCCACTTTTTAGCA<br>ATATGATAAATACCCAATTTGATAAATTTATCTTGCCTCATAGTTATCGTAAAGATAAACTTTACCAAAA<br>TGATAAATTGAGGTGTGGGTATGACTAAAAAAAATTTCATTCATTATTAACGGCCAG |
| JSO-1758: pTrc backbone | GTTTTGGCGGATGAGAGAAG |
| JSO-1759: pTrc backbone | GGTTTATTCCTCCTTATTTAATCGATACATTAA |
| JSO-1756: *clbS* (into pTrc) | ATGGCTGTTCCATCATCAAAAGAAGAG |
| JSO-1757: *clbS* (into pTrc) | CTATTCTGCAAGACATTTCTGCAGTTTATTTAACC |
| JSO-1766: *clbS* deletion from pTrc-*clbS* (construction of pTrc-Δ*clbS*) | TTCTTTTGATGATGGAACAGCCAT |
| JSO-1767: *clbS* deletion from pTrc-*clbS* (construction of pTrc-Δ*clbS*) | CTGCAGAAATGTCTTGCAGAATAG |
| JSO-1727: $P_R$-*lux* insert | TGAATGAAATTTTTTTAGTCATACAACCTCCTTAGTACATGCA |
| JSO-1730: $P_R$-*lux* insert | GGTACCGCTCGAGTTAATTAAGATCAGCCAAACGTCTCTTC |
| JSO-1691: $P_R$-*lux*, $P_{metako}$-*lux* and pEVS143-empty (kanR) | TTAATTAACTCGAGCGGTACCCG |
| JSO-1692: $P_R$-*lux* and $P_{metako}$-*lux* | ATGACTAAAAAAATTTCATTCATTATTAACGGCCAG |
| JSO-1731: pBR322-empty (*ampR*) | TTAATTAACTCGAGCGGTACCTCTTAC |
| JSO-1743: pBR322-empty (*ampR*) and pEVS143-empty (*kanR*, with JSO-1691) | GGATCCGGTGATTGATTGAGC |
| JSO-1741: empty BAC2 (cos sites removed) | GTATTTTGTCCACATAACCGTGCG |
| JSO-1742: empty BAC2 (cos sites removed) and cmR (with JSO-1745) for pEVS143-empty (*kanR-cmR*, ligated with primer product used for pEVS143-empty *kanR*) | GAGTGAGCTAACTCACATTAATTGCG |
| JSO-1745: *cmR* for pEVS143-empty (*kanR-cmR*); ligated with primer product used for pEVS143-empty (*kanR*) | GGCATTTATTCTCAGGATAATTGTTTCAGC |
| JWO-1046: GC-rich probe | GCCGCATGCCGCATGCCGCATGCCGCATGCCGCATGCCGCATGCCGCATGCCGCATGCCGCAT |
| JWO-1047: GC-rich complement | ATGCGGCATGCGGCATGCGGCATGCGGCATGCGGCATGCGGCATGCGGCATGCGGCATGCGGC |
| JWO-1044: AT-rich probe | CACACATTTGCAAATTTCGCAAATTTGCACAGATCTTGCAAATTTGCAAATTTGCAAATTTCA |
| JWO-1045: AT-rich complement | TGAAATTTGCAAATTTGCAAATTTGCAAGATCTGTGCAAATTTGCGAAATTTGCAAATGTGTG |

### Supplementary References

1. Baba, T. *et al.* Construction of Escherichia coli K-12 in-frame, single-gene knockout mutants: the Keio collection. *Mol. Syst. Biol.* **2**, 2006.0008 (2006).
2. Eames, M. & Kortemme, T. Cost-Benefit Tradeoffs in Engineered lac Operons. *Science* **336**, 911–915 (2012).
3. Tomkovich, S. *et al.* Locoregional Effects of Microbiota in a Preclinical Model of Colon Carcinogenesis. *Cancer Res.* **77**, 2620–2632 (2017).
4. Owen, S. V. *et al.* Characterization of the Prophage Repertoire of African Salmonella Typhimurium ST313 Reveals High Levels of Spontaneous Induction of Novel Phage BTP1. *Front. Microbiol.* **8**, (2017).
5. Owen, S. V. *et al.* Prophage-encoded phage defense proteins with cognate self-immunity. *bioRxiv* 2020.07.13.199331 (2021) doi:10.1101/2020.07.13.199331.
6. Novick, R. Properties of a cryptic high-frequency transducing phage in Staphylococcus aureus. *Virology* **33**, 155–166 (1967).
7. Ubeda, C., Barry, P., Penades, J. R. & Novick, R. P. A pathogenicity island replicon in Staphylococcus aureus replicates as an unstable plasmid. *Proc. Natl. Acad. Sci.* **104**, 14182–14188 (2007).
